# tRNA Modification Landscapes in Streptococci: Shared Losses and Clade-Specific Adaptations

**DOI:** 10.1101/2025.10.13.681863

**Authors:** Ho-Ching Tiffany Tsui, Chi-Kong Chan, Yifeng Yuan, Roba Elias, Jingjing Sun, Virginie Marchand, Marshall Jaroch, Guangxin Sun, Irfan Manzoor, Ana Kutchuashvili, Yuri Motorin, Grazyna Leszczynska, Kinda Seaton, Kelly C. Rice, Manal A. Swairjo, Malcolm E. Winkler, Peter C. Dedon, Valérie de Crécy-Lagard

**Author notes:** Current address: Kihealth Inc., Saint Augustine, Florida, USA.

## Abstract

tRNA modifications are central to bacterial translational control. Here, we integrated genetics, mass spectrometry, epitranscriptomics, and comparative genomics to map the tRNA modification genes of the Gram-positive pathogens *Streptococcus mutans* and *Streptococcus pneumoniae*. Both species show a marked loss of modifications dependent on Fe–S enzymes, consistent with a broader trend of Fe–S enzyme reduction in Streptococcus central metabolism. In addition, the D, m^1^A, m^7^G, t^6^A, and i^6^A modifications were mapped in *S. pneumoniae* tRNAs, and we confirmed that a unique DusB1 enzyme is responsible for the insertion of all the detectable D modifications. We uncovered differences in queuosine (Q) metabolism: while *S. mutans* synthesizes Q de novo, *S. pneumoniae* instead salvages preQ₁ and accumulates the epoxy-Q precursor, a strategy shared with multiple other Streptococci as revealed by analysis of Q pathways in 1,599 sequenced streptococcal genomes. Comparative essentiality profiling of modification genes revealed notable differences, including the essentiality of the NLJ-threonylcarbamoyladenosine (tLJA) synthesis enzyme TsaE in *S. pneumoniae* but not in *S. mutans*, which was confirmed by genetic studies. We found that suppressor mutations in *asnS* encoding asparaginyl-tRNA synthetase (AsnRS) restored viability to Δ*tsaE* mutants, albeit with reduced growth. Our finding highlights the functional importance of modifications in the recognition of tRNAs by aminoacyl-tRNA synthetases.

## Introduction

Streptococci are a diverse group of Gram-positive bacteria that are common members of the human microbiota, capable of colonizing the skin as well as mucosal membranes of the oral cavity, upper respiratory tract, intestines, and vaginal tract [1]. Clinically important species include *Streptococcus pyogenes* (Group A), responsible for pharyngitis, rheumatic fever, and necrotizing fasciitis[2]; *Streptococcus agalactiae* (Group B), a cause of neonatal infections [3]; *Streptococcus mutans,* a keystone pathogen in the development of dental caries, as well as infective endocarditis in at-risk patient groups [4]; and *Streptococcus pneumoniae*, a leading cause of otitis, pneumonia and meningitis [5]. These bacteria exhibit a wide range of virulence factors and adaptation mechanisms, making them important subjects of study in both basic microbiology and infectious disease research.

Transfer RNAs (tRNAs) are the essential decoding molecules in the translation process, and they undergo extensive post-transcriptional modification. These modifications fine-tune translation efficiency, fidelity, and tRNA stability [6,7]. In pathogenic bacteria, tRNA modifications can modulate stress responses [8,9], antibiotic resistance [9,10], metabolic fluxes [11] and the accurate synthesis of infection-related proteins [12,13] and are hence important in microbial pathogenesis and virulence in many different bacteria (see [14–16] for recent reviews). In addition, a few bacterial tRNA modification enzymes are essential and have no ortholog in humans and are thus being explored as antibacterial targets [17,18].

That said, our understanding of the role of tRNA modifications in pathogens remains limited, as comprehensive mapping of specific tRNA modifications and their corresponding biosynthetic enzymes have been completed in only a few organisms, including the model gram-negative *Escherichia coli* K12 [17]. For decades, the intracellular pathogen *Mycoplasma capricolum* was the only Gram-positive organism with combined experimental and bioinformatic evidence linking tRNA modifications with their genes [17,19]. As discussed previously [17], no single method, analytical or computational, can accurately predict the full set of tRNA modification enzymes in a given organism. However, the combination of liquid chromatography-tandem mass spectrometry (LC-MS/MS), tRNA sequencing, and bioinformatics methods greatly increases the accuracy of the predictions [17]. This type of integrative approach has enabled the mapping of modifications and their genes in three additional gram-positive bacteria: *Bacillus subtilis [20]*, *Mycobacterium tuberculosis* [21] and *Streptomyces albidoflavus* [22]. The initial analysis of tRNA modification genes in *B. subtilis* revealed open questions [20], including the specificities of dihydrouridine synthase enzymes and the identity of the enzymes involved in cmnm^5^(s^2^)U deacetylation and nm^5^(s^2^)U methylation that have now both been resolved [23,24](**Fig. 1, Supplemental Data 1A**).

**Figure 1.**
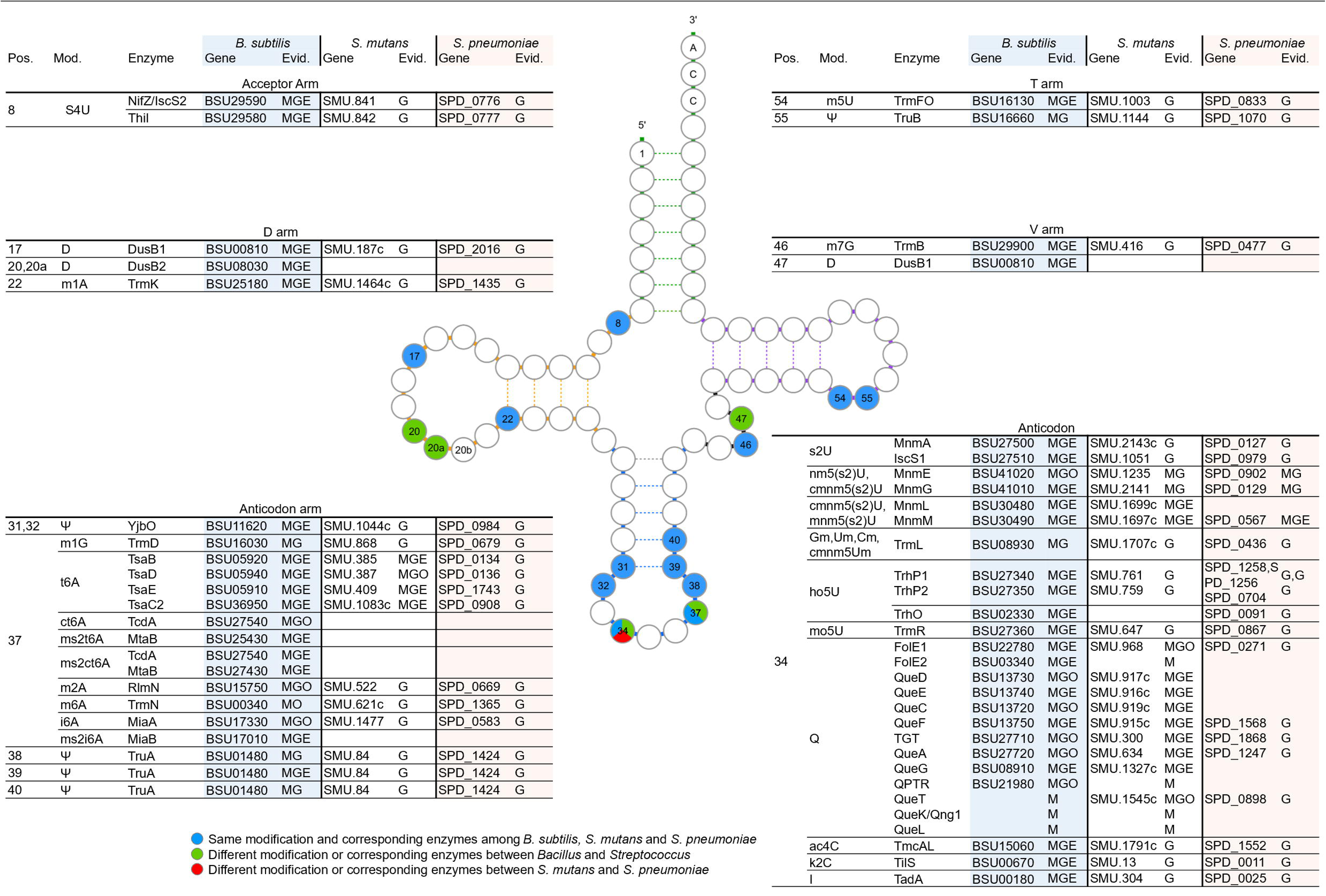
The comparison of tRNA modifications and corresponding enzymes among *B. subtilis*, *S. mutans*, and *S. pneumoniae*. Modified positions are colored blue when the modification and enzymes are conserved and orthologous in three strains; green when they differ between *Bacillus* and *Streptococcus;* and red when they differ between *S. mutans* and *S. pneumoniae*. Evidence code: M, modification reported; G, gene coding modifying protein or an ortholog of an existing modifying protein; E, experiments that validated modifying gene(s) of the species via in vivo gene knockout or in vitro expression; O, ortholog that was validated experimentally in a different species. References for all proteins listed are given in Supplemental data S1D.

Dihydrouridine (D) is a widespread modification that is important for tRNA structure and flexibility [25]. Three subfamilies of dihydrouridine synthases with different specificities modify *E. coli* tRNAs. DusA (K05539) modifies positions 20 and 20a, DusB (K05540) position 17, and DusC (K05541) position 16. Only two DusB paralogs are present in *B. subtilis,* DusB1 (BSU00810) and DusB2 (BSU08030), and it was recently shown that in this model organism, DusB1 exhibits multisite enzyme activity, enabling D formation at positions 17, 20, 20a, and 47, while DusB2 specifically catalyzes U to D conversion at positions 20 and 20a [23]. Many pathogenic Bacillota (formerly Firmicutes) have lost one of the two DusB enzymes [26]. Many *Staphylococcus aureus* strains encode a full-length DusB2 and a truncated version DusB2-C, but only the full-length one is active [27], while *M. capricolum* and most *Streptococci* only encode a DusB1 [23,26]. Heterologous expression studies with the Mycoplasma DusB1 (MCAP_0837) enzyme showed that it modifies positions 17, 20, and 20a, but this is yet to be validated genetically in an endogenous host [26].

Mnm^5^s^2^U is a common wobble base (position 34) modification found in both Gram^+^ and Gram^-^ bacteria that is important for the decoding of NNA/NNG codons, particularly in split codon boxes [7]. The MnmEG complex catalyzes the first step of this complex pathway both in *B. subtilis* and *E. coli* [28,29] producing the cmnm^5^s^2^U or nm^5^s^2^U precursors. Several studies have linked the absence of MnmG (or MnmE) with pleiotropic phenotypes in many bacteria [30] including virulence defects in different Streptococci [31–34]. The subsequent stages of the pathway are catalyzed by non-orthologous enzymes in the two major model organisms. The *E. coli* FAD-dependent cmnm^5^(s^2^)U deacetylase [MnmC1/MnmC(o), a domain of the bifunctional MnmC (K15461) protein] is replaced in *B. subtilis* by the radical SAM enzyme MnmL (K07139)[24]. Two different families of methylases finalize the synthesis of mnm^5^U by methylating nm^5^U: the *E. coli* MnmC2/MnmD/MnmC(m) type and the *B. subtilis* MnmM type [24,35]. The functional roles of the *S. mutans* MnmL and MnmM orthologs have also been confirmed by genetic studies [24]. The corresponding genes do accumulate inactivating mutations in many *S. pneumoniae* strains that lack mnm^5^s^2^U and accumulate the nm^5^s^2^U precursor [24]. In *S. mutans,* the only other tRNA modification genes that have been characterized are involved in the synthesis of two other complex modifications of the tRNA Anticodon-Stem-loop (ASL): the wobble base modification Queuosine (Q)[36] at position 34 and the universal N6 adenosine modification-threonylcarbamoyladenosine (t^6^A) at position 37 [37].

Q is a deazapurine derivative synthesized de novo from GTP in many bacteria in seven catalytic steps [38]. Q precursors such as preQ_0_, preQ_1,_ or the queuine base (q) can be salvaged in some bacteria [38] using a wide variety of transporters, including the QPTR and QueT/QtrT subgroups of the Energy-coupling factor (ECF) transporter families [38,39]. Pathogenic bacteria that colonize the human host can use salvaged q either directly by mutating the substrate specificity of the bacterial TGT enzyme from preQ_1_ to q [40], or indirectly by converting salvaged q to preQ_1_, a reaction catalyzed by a radical-SAM enzyme QueL [40]. Queuine can be generated by cleaving Q with hydrolases such as QueK which has also been characterized in a human pathogen [40]. S*. mutans* can synthesize Q *de novo* using the same pathway as *E. coli,* and all Q synthesis genes have been experimentally validated, including the signature enzyme TGT that inserts the preQ_1_ precursor into target tRNAs [36]. In addition, we confirmed *S. mutans* can salvage preQ_0_ and preQ_1_ using a QueT-dependent ECF transporter [36]. No obvious phenotype was linked to Q absence, but it was found that Q levels in *S. mutans* tRNAs were greatly influenced by media conditions [36]. It remains to be determined whether *S. mutans* or any other Streptococci can recycle or salvage q or Q.

t^6^A is a universal modification and its synthesis requires two steps, catalyzed in bacteria by TsaC (or TsaC2) and the TsaBDE complex, respectively [41,42]. The essentiality of the t^6^A synthesis genes varies among organisms. They are essential in *E. coli* and *S. aureus* but not in *Deinococcus radiodurans*, *B. subtilis,* or *S. mutans* [43]. In some organisms, t^6^A is a determinant for the charging of tRNA^Ile^ by Isoleucyl-tRNA synthetase [41,43] and could also be a determinant for lysidine synthase (TilS), which is one of the rare essential tRNA modification enzymes, as it is required for cognate tRNAs to decode AUA (Ile) codons [43]. However, it seems many organisms can circumvent these t^6^A requirements by mechanisms that have not been fully elucidated.

In this work, we combined analytical, genomic, genetic, and bioinformatic approaches to predict the complete sets of tRNA modification enzymes in two model Streptococci (*S. mutans* and *S. pneumoniae)*. Our analysis revealed shared differences with the Gram-positive model *B. subtilis*, notably a consistent loss of iron–sulfur cluster–dependent modifications. We also found that the Q pathway is markedly reduced in *S. pneumoniae* compared to *S. mutans*, a pattern representative of most sequenced Streptococci, as shown by our comparative analysis of ∼1,600 genomes. Finally, we demonstrate that, unlike in *S. mutans*, where t^6^A is dispensable, this conserved anticodon–stem–loop modification is essential in *S. pneumoniae*, although suppressor mutations arise at high frequency, pointing to previously unrecognized roles of t^6^A in translation.

## Methods

### Databases resources

Different databases and repositories from the NCBI resources (https://www.ncbi.nlm.nih.gov/) [44] including Pubmed (https://pubmed.ncbi.nlm.nih.gov/) and RefSeq (https://www.ncbi.nlm.nih.gov/refseq/) [45] and COG (https://www.ncbi.nlm.nih.gov/research/cog/) [46], from the EBI resources (https://www.ebi.ac.uk/) [47] including Uniprot (https://www.uniprot.org/) [48] and InterPro (https://www.ebi.ac.uk/interpro/) [49], from the KEGG resources (https://www.genome.jp/kegg/) [50] including the KO (KEGG ORTHOLOGY) (https://www.genome.jp/kegg/ko.html) database, the Protein Data Bank (http://www.rcsb.org/) [51] and the BV-BRC (https://www.bv-brc.org/) [52] resources were routinely used. Subtiwiki (https://subtiwiki.uni-goettingen.de/v5/welcome) was used as the source of rRNA modification genes in Gram-positive [53].

### Q synthesis and salvage gene/protein analyses

Genome DNA sequences and protein sequences from 1,599 *Streptococcus* strains were retrieved from the NCBI database in July 2025, applying filters for annotation (‘RefSeq’) and assembly status (‘complete’) (**Supplemental data 1F** and **Fig. S1 step 4**). Q proteins and Q genes were searched in them using BLASTp and tBLASTn [54] with a P-value cutoff of 1E-10 and an identity cutoff of 20% (**Fig. S1 step 5**). Query sequences were the Queuosine synthesis and salvage proteins from *B. subtilis* **(Supplemental Data 1A and 1E)**. Pseudogenes were called when matches were found by tBLASTn but not by BLASTp, indicative of internal stops or frameshifts. A hundred and twenty conserved marker proteins were identified and aligned using GTDBtk (v2.3.2) [55] with *Pseudolactococcus* and *Lactococcus* as the outgroup. A maximum likelihood tree of the concatenated marker proteins was generated using IQ-TREE2 (v2.2.2.7) [56], employing the LG+G substitution model and 1,000 fast bootstrap replicates (**Fig. S1 step 6**). The tree was visualized with the mapping of Q protein patterns using iTOL (v7.2.1) [57]. Branches of the same species harboring the same Q proteins were collapsed and represented by triangles, the size of which correlates with the number of species. Species names were updated when their RefSeq classification disagree with their GTDB classification or their position in the tree (**Fig. S1 step 7**, **Supplemental Data 1E**). Species represented by fewer than three strains or rare Q gene patterns were hidden. [Multiple sequence alignment of Q proteins (input sequences in **Supplemental Data 1F**) was performed using MAFFT (v7.520) [58] with setting “--maxiterate 1000 --localpair”. Sequence logos were generated with the aligned sequences using Weblogo [59]. Alignments of genome sequences was performed using BLASTn (v2.15).

### Sequence similarity network analyses

The sequence similarity network (SSN) of inosine/uridine-preferring nucleoside hydrolase (IPR023186) family was generated using EFI-EST (EFI Enzyme Similarity Tool, efi.igb.illinois.edu/efi-est)[60](**Fig. S1 step 8**). 36,956 sequences in the family were retrieved from UniProt and subjected to EFI-EST using the family option. Each node in the network represents one or multiple sequences that share no less than 90% identity. The initial SSN was generated with an Alignment Score Threshold (AST) set such that each connection (edge) represented a sequence identity above 40%. The nodes from *Streptococcus* were colored and visualized using Cytoscape 3.10.1[61]. More SSNs were created by gradually increasing the alignment score cutoff in small increments (usually by 5 AST). This process was repeated until paralogs in the IPR023186 family were separated into different clusters (AST=100). In the final view of the SSN, the shapes of nodes are based on the kingdom of species of the representative sequence of each node. The nodes of *Streptococcus* are filled in yellow. The nodes that have been annotated experimentally as not being QueK are filled in black. The nodes are filled in blue and skyblue when their encoding gene is next to *queT* or other transporter gene, respectively. The borders of nodes are highlighted in red when the representative sequences of each node are from genomes that lack *queDECF* genes. For better visualization, clusters with less than 30 nodes are hidden. The resulting SSN was then subjected to EFI-EST and EFI-GNT for gene neighborhood analysis with default settings. The retrieved genome neighborhood was visualized using Gene Graphics (https://genegraphics.net/)[62]. Identifiers for sequences used in the SSN and in the genome neighborhood analyses are available in **Supplemental Data S1G and S1H**.

### Codon usage analyses

The genome distances between each of the 234 *S. pneumoniae* genomes and all other Streptococci present in our genome set were calculated using MASH v2.3[63] (**Fig. S1 step 9**).

26 genome pairs that each share at least 92% genome identity (distance < 0.08) between a *S. pneumoniae* strain and a member from another Streptoccocal clade were chosen (**Supplemental Data S1I**). The coding DNA sequences (CDSs) of the selected genome pairs were retrieved from the NCBI database. After removing pseudogenes, the number of each of the 61 sense codons was calculated per CDS (excluding start codons) (**Fig. S1 step 10**). Then, the genome-wide codon usage for every amino acid was compared using a paired two-tailed t-test (**Supplemental Data S1I**).

## Structural analysis

Experimental structures of AsnRS, AspRS and LysRS were obtained from the Protein Data Bank (https://www.rcsb.org/), and the AlphaFold models for *Sp*AsnRS and *Sm*AsnRS were obtained from UniProt (IDs Q8CWQ4 and Q8DTM2, respectively). Structures were aligned and analyzed in Pymol (https://www.pymol.org/) [64]. The structure-based multi-sequence alignment was generated using the PROMALS3D online tool (http://prodata.swmed.edu/promals3d/)[65] and printed in ESPript [66].

### Strains and culture conditions

All bacterial strains used in this work are listed in **Table S1**. Strains of *S. mutans* were routinely cultured statically at 37°C in 5% CO_2_ in BHI medium (BD Biosciences) with 10 µg/mL erythromycin (Sigma) or 1 mg/mL kanamycin (Sigma) when appropriate. *S. pneumoniae* derivatives were derived from unencapsulated strains IU1824 (D39 Δ*cps rpsL1*) and IU1945 (D39 Δ*cps*), which were derived from the encapsulated serotype-2 D39W progenitor strain IU1690 [67,68]. Strains containing antibiotic markers were constructed by transformation of CSP1-induced competent pneumococcal cells with linear DNA amplicons synthesized by overlapping fusion PCR [69]. Strains containing markerless alleles in native chromosomal loci were constructed using allele replacement via the P_c_-[*kan*-*rpsL*^+^] (Janus cassette)[70]. Primers and DNA templates used to synthesize different amplicons are listed in **Table S2**. *S. pneumoniae* were grown on plates containing trypticase soy agar II (modified; Becton-Dickinson), and 5% (vol/vol) defibrinated sheep blood (TSAII-BA). Plates were incubated at 37°C in an atmosphere of 5% CO_2_. TSAII-BA plates for selections contained antibiotics at concentrations described previously [69]. Bacteria were cultured statically in Becton-Dickinson brain heart infusion (BHI) broth at 37°C in an atmosphere of 5% CO_2_, and growth was monitored by OD_620_ as described before [69]. Mutant constructs were confirmed by PCR and DNA sequencing of chromosomal regions corresponding to the amplicon region used for transformation. Ectopic expression of various genes was achieved with a P_Zn_ zinc-inducible promoter in the ectopic *bga*A site. 1/10 concentration of Mn^2+^ was added with Zn^2+^ to prevent zinc toxicity [69]. 0.1 to 0.2 mM (Zn^2+^/(1/10) Mn^2+^) was added to TSAII-BA plates or BHI broth for inducing conditions. Mn^2+^ was added with Zn^2+^ to prevent zinc toxicity [69,71]. Ectopic expression of *asnS* [D121A] was achieved with a constitutive promoter P*_ftsA_* at the *bgaA* site [72].

In all experiments, *S. pneumoniae* cells were inoculated from frozen glycerol stocks into BHI broth, serially diluted, and incubated 12–15 h statically at 37°C in an atmosphere of 5% CO_2_. For culturing merodiploid strains IU18963 (Δ*tsaE*//P_Zn_-*tsaE*^+^) that require Zn^2+^ for expressing *tsaE* from a Zn-dependent promoter (P_Zn_), 0.1 mM (Zn^2+^/(1/10) Mn^2+^) was added to BHI broth in the overnight cultures. The next day, cultures at OD_620_ ≈0.1–0.4 were diluted to OD_620_ ≈0.003 in BHI broth with no additional (Zn^2+^/(1/10) Mn^2+^) or the amounts of (Zn^2+^/(1/10) Mn^2+^) indicated for each experiment. Doubling time determination was performed by first examining the growth curves on a log scale to determine the time points when growth was in the exponential phase. Doubling times were determined with the exponential growth equation using only data points that exhibit exponential growth using GraphPad Prism version 10.0.0 for Windows, GraphPad Software, Boston, Massachusetts USA, www.graphpad.com. Maximal growth yields were determined by the highest OD_620_ values obtained within 9 h of growth. Cultures were sampled for phase microscopy at early to mid-exponential phase.

### *S. pneumoniae* Tn**-**seq transposon library generation and insertion sequencing

Tn-seq was carried out according to protocols described in [73] using a transposon insertion library generated for WT D39 Δ*cps rpsL1* (IU1824) and WT D39 *cps^+^ rpsL1* (IU1781) cultured with BHI media. In brief, transposon library starter cultures were thawed, and were diluted to OD_620_LJ≈LJ0.005 in 5 ml of BHI and were grown at 37°C with 5% CO_2_ to OD_620_LJ≈LJ0.4. 5LJml of culture at OD_620_LJ≈LJ0.4 were used to extract genomic DNA. *MmeI* digested DNA were ligated to adaptors and sequenced on the Illumina NextSeq 500 at the Center for Genomics and Bioinformatics, Indiana University Bloomington. Data were mapped and analyzed, and Tn insertion data were visualized graphically using the Artemis genome browser (version 10.2) [74].

### *S. pneumoniae* transformation assays

Transformations were performed as previously described [72,75]. Δ*tsaE*::P_c_-*erm,* Δ*tsaE* P*_c_-kan-rpsL* amplicons, and positive control Δ*bgaA*::P_c_-*erm* amplicon were synthesized by PCR using the primers and templates listed in Table S2 and contain ≈1 kb of flanking chromosomal DNA. A fusion Δ*tsaE* markerless amplicon or an amplicon from strain IU18963 were used for transformation of IU18886. All transformation experiments were performed using no added DNA as the negative control. The sizes of colonies indicated in **Table 1** were relative to colonies transformed with the WT strain with a control Δ*bgaA* amplicon. For transformations in 0.1 mM or 0.2 mM (Zn^2+^/(1/10) Mn^2+^), ZnCl_2_ and MnSO_4_ stock solutions were added to transformation mixes and soft agar for plating and spread onto blood plates containing the respective (Zn^2+^/(1/10) Mn^2+^) concentrations to induce gene expression under control of the P_Zn_ zinc-inducible promoter in the ectopic *bgaA* site.

**Table 1.**
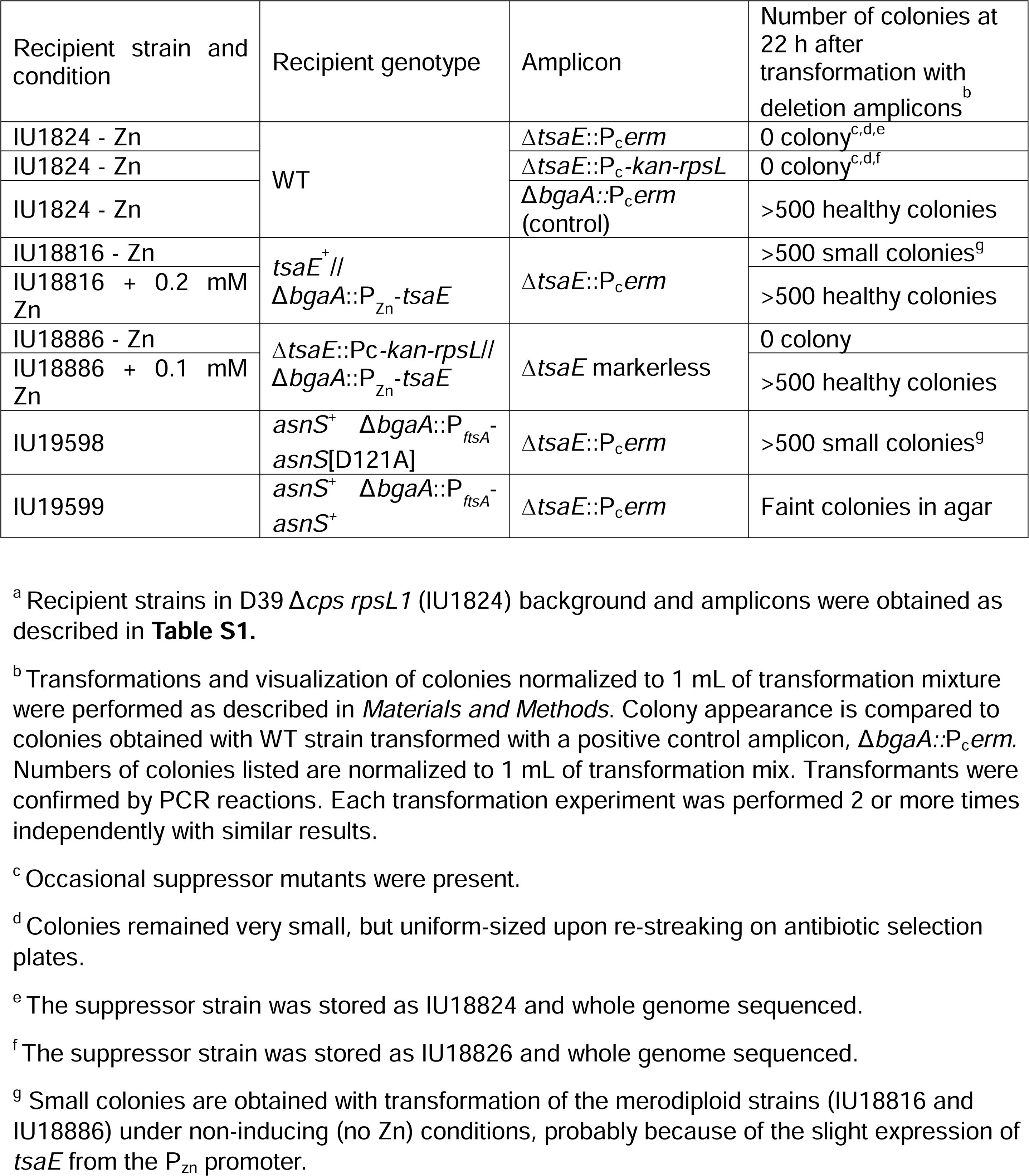
Transformation of Δ*tsaE* amplicons into different backgrounds confirms *tsaE* essentiality in *S. pneumoniae* D39 and suppression by *asnS*[D121A] mutation^a^.

### Whole-genome DNA sequencing

Whole-genome sequencing was employed to identify suppressor mutations and verify the genomes of the constructed mutants. Strains IU18824, IU18826, E797, and K797 (**Table S2**) containing suppressor mutations that allowed growth of a Δ*tsaE* mutant were isolated as described in Results. Genomic DNA preparation, DNA library construction, Illumina MiSeq DNA sequencing, and bioinformatics analyses were performed as described previously [75]. Genomic DNA from the *S. mutans tsaE* deletion strain (JB409) and its isogenic WT strains (*S. mutans* UA159, Wen laboratory Isolate) were extracted using DNeasy Blood and tissue kit (Qiagen), following the manufacturer’s instructions. and was sequenced at the Microbial Genome Sequencing Center (MiGS) Illumina sequencing libraries were prepared using the tagmentation-based and PCR-based Illumina DNA Prep kit and custom IDT 10bp unique dual indices (UDI) with a target insert size of 280 bp. No additional DNA fragmentation or size selection steps were performed. Illumina sequencing was performed on an Illumina NovaSeq X Plus sequencer in one or more multiplexed shared-flow-cell runs, producing 2×151bp paired-end reads. Demultiplexing, quality control and adapter trimming were performed with bcl-convert1 (v4.2.4), a proprietary Illumina software for the conversion of bcl files to basecalls. Illumina generated 2×151bp paired end read data was used as the input for variant calling against the reference (GenBank id: AE014133) Variant calling was carried out using BreSeq1 under default settings [76].

### tRNA and rRNA extraction

Bulk tRNA samples from *S. mutans* cells were prepared in previous studies [24,36]. tRNA samples from *S. pneumoniae* derivatives were prepared as described in detail previously (see MW methods in [24]). Overnight cultures (12–15 h) diluted to an initial OD_620nm_ 0.005 and allowed to grow until an OD_620nm_ of ≈ 0.2. For cultures treated with PreQ_0_ (Millipore Sigma, AMBH2D6F0DFD) or PreQ_1_ (Millipore Sigma, product # SML0807), stock solution of 100 µM PreQ_0_ or PreQ_1_ in DMSO was added to BHI media to a final concentration of 100 nM at initial growth. The same volume of DMSO was added to the control samples. The purified tRNA samples were stored at −80°C until further analysis.

For rRNA samples, the WT *Streptococcus mutans* UA159 strain was cultured in BHI broth at 37°C in 5% CO₂ until mid-log phase (OD₆₀₀= 0.6–0.8). Cells were harvested and lysed as described by Chia et al. [77], followed by TRIzol extraction (Invitrogen) and purification using the NucleoBond RNA/DNA kit (Macherey-Nagel) according to the manufacturer’s instructions. The resulting preparation was treated with DNase I (NEB) for 10 minutes at 37 °C, and the RNA was then cleaned up using the RNeasy Mini Kit (Qiagen). RNA concentration and purity were assessed by NanoDrop spectrophotometry, and integrity was verified by running a 1 % agarose; The RNA electropherogram and RNA Integrity Number (RIN) were obtained using an Agilent 4200 TapeStation (ICBR, UF).

### Phase microscopy and cell measurements

Cultures were sampled for phase microscopy at OD_620_ ≈0.1–0.2 for IU1824 and IU18963, and OD_620_≈0.03–0.05 for IU18824. 500 µL of culture were centrifuged at 20,000g at room temperature for 3 min. 450 µL of supernatant was removed and the pellet was resuspended in the remaining 50 µL media. 1.5 µL of cells were placed on a slide and covered with a coverslip and observed with phase contrast microscopy using a Nikon Eclipse E400 microscope.

Cell lengths and widths of strains growing in BHI broth were measured from phase-contrast images by using Nikon NIS-Element BR software as described before [69]. Only ovoid-shape predivisional cells were measured. Cells (88 or more) from 2 independent experiments were measured and plotted with scatter plot. P values were obtained by one-way ANOVA analysis by using the nonparametric Kruskal-Wallis test in GraphPad Prism program.

### RNA hydrolysis for LC-MS/MS analysis

Purified tRNA and rRNA (3 µg) were hydrolyzed to nucleosides as previously described [78] in a 50-µL enzyme cocktail containing 8 U benzonase (Sigma), 5 U calf intestinal alkaline phosphatase (Sigma), 0.15 U snake venom phosphodiesterase I (Sigma), 5 ng coformycin NCI), 0.1 mM deferoxamine (Sigma), 0.1 mM butylated hydroxytoluene (Sigma), 50 nM internal standard [^15^N]_5_-deoxyadenosine (Cambridge Isotope Laboratories), 2.5 mM MgCl_2_ (Sigma), and 5 mM Tris-HCl buffer pH 8.0 (Invitrogen). Enzyme control reactions were prepared identically but without the addition of RNAs. The reaction mixtures were gently tapped, briefly spun down, and incubated at 37 °C for 6 h. After hydrolysis, the samples were centrifuged at 3,000 × *g* at 4°C for 10 min and analyzed by LC-MS/MS.

### LC-MS/MS analysis of modified ribonucleosides

The LC-MS/MS analysis of tRNAs extracted from *S. mutans* was described previously [24,36]. For *S. pneumoniae*, a first batch was analyzed as described in [24]. All other LC-MS/MS analyses were performed on an Agilent 1290 Infinity II UHPLC system equipped with an inline diode array detector (DAD) and coupled to an Agilent 6495c triple quadrupole mass spectrometer (MS). For each run, 180 ng of hydrolyzed RNA was injected onto a Waters ACQUITY UPLC BEH C18 column (50 × 2.1 mm i.d., 1.7 µm) equipped with an ACQUITY UPLC BEH C18 VanGuard Pre-column (5 × 2.1 mm i.d., 1.7 µm) and an ACQUITY Column In-line Filter (0.2 µm). To facilitate the detection of low abundance modified ribonucleosides, a second injection of 600 ng of hydrolyzed RNA was performed for each sample. The column was maintained at 30 °C and operated at a flow rate of 0.3 mL/min with mobile phase solvents consisting of Buffer A (0.02% formic acid in water) and Buffer B (0.02% formic acid in 70% aqueous acetonitrile). The gradient of Buffer B was programmed as follows: 0–5 min, 0–1%; 5–7 min, 1–3%; 7–9 min, 3–7%; 9–10 min, 7–10%; 10–12 min, 10–12%; 12–13 min, 12–15%; 13–15 min, 15–20%; 15–16 min, 20–75%; 16–17 min, 75–100%; 17–20 min, held at 100%, 20–21 min, 100–0%; and 21–30 min, re-equilibrated at 0%. The DAD wavelength was set to 260 nm to acquire signals of canonical ribonucleosides. The JetSream ESI source operated in positive-ion mode with the following optimized parameters: drying gas temperature, 200 °C; gas flow, 11 L/min; nebulizer, 20 psi; sheath gas temperature, 300 °C; sheath gas flow, 12 L/min; capillary voltage, 3000 V; and nozzle voltage, 0 V. MS/MS analysis leveraged a dynamic multiple reaction monitoring (dMRM) mode with retention time windows set as ± 2 min and collision energies (CEs) optimized for maximal sensitivity. Retention time of modified ribonucleosides were confirmed with synthetic standards (**Supplemental Data 2A**) and served as a primary criterion for peak identification. Raw peak areas for each modified ribonucleoside were extracted using Agilent QQQ Quantitative Analysis version 10.2 and normalized to the sum of the peak areas for the four canonical ribonucleosides recorded by the DAD. Raw LC-MS/MS data have been deposited to the ProteomeXchange Consortium (http://proteomecentral.proteomexchange.org) via the PRIDE (http://www.ebi.ac.uk/pride/) partner repository with the dataset identifier PXD069173.

Besides the modified ribonucleosides for which we have synthetic standards, we also monitored a set of putative modifications that lack available synthetic standards in our inventory but have been previously reported in other prokaryotes. Information regarding precursor and product ions for these modifications was obtained from Modomics [79], precedent studies, and the literature, and incorporated into the dMRM method. Identities for these putative modifications were validated using a two-step approach. First, extracted ion chromatograms from the unit-resolution LC-MS/MS analysis of hydrolyzed tRNAs were compared with those from enzyme control reactions; only peaks observed exclusively in the hydrolyzed tRNAs were considered for further analysis. These peaks were subsequently subjected to high-resolution LC-MS analysis using a Thermo Fisher Scientific Dionex Ultimate 3000 UHPLC system coupled to an Orbitrap Q Exactive mass spectrometer equipped with an Ion Max source and a heated ESI (HESI II) sprayer. For each injection, 2 µg of hydrolyzed tRNAs was loaded onto the same analytical column, eluted with the same mobile phase solvents, and subjected to the same LC gradient and column oven temperature as described above. The source and MS parameters are as follows: spray voltage, +4.2 kV; capillary temperature, 320 °C; sheath gas, 50 arbitrary unit (au); auxiliary gas, 15 au; spare gas, 3 au; max spray current, 100 µA; probe heater temperature, 400 °C; and S-Lens RF level, 70 au. Initially, the hydrolyzed tRNAs were analyzed using a targeted selected ion monitoring (SIM) mode with a mass inclusion list of putative modifications identified from the LC-MS/MS analysis. The Automatic Gain Control (AGC) target was set at 20,000, with a maximum injection time of 150 ms. SIM mass spectra were acquired at a resolution of 35,000 full width at half maximum (FWHM) at m/z 200 with an isolation window of 4.0 Da. Peaks with a mass error < 5 ppm were subjected to targeted MS/MS fragmentation. To this end, the AGC target was increased to 500,000, with a maximum injection time of 100LJms. MS/MS spectra were acquired were acquired over an *m/z* range of 50 – 450 at a resolution of 17,500 FWHM at *m/z* 200 with an isolation window of 2.0 Da and CEs between 10 and 20 eV. The acquired spectra were compared to those reported in literature or to fragmentation patterns predicted using the Mass Fragmentation Tool in ChemDraw version 22.0.

### Analysis of tRNA modifications by AlkAnilineSeq

tRNA samples (∼200 ng) from *S. mutans* and *S. pneumoniae* were subjected to random fragmentation by alkaline hydrolysis in 50 mM sodium-bicarbonate buffer at pH 9.2 and 96°C for 5 min [80]. The reaction was stopped by ethanol precipitation using 3M Na-OAc, pH 5.2 and glycoblue. After centrifugation, the RNA pellet was washed with 80% ethanol and resuspended in nuclease-free water. RNA fragments were de-phosphorylated by Antarctic phosphatase (NEB ref M0289L, USA) at 37°C for 1h and precipitated using 3M Na-OAc, pH 5.2 and glycoblue as previously described [81]. After centrifugation, the RNA pellet was washed with 80% ethanol and the pellet was resuspended in 1M Aniline pH 4.5 and incubated for 15 min at 60°C in the dark [82] The reaction was stopped by ethanol precipitation using 3M Na-OAc, pH 5.2 and glycoblue. The pellet was washed twice with 80% ethanol, dried and resuspended in 3.5 µL of nuclease free water. RNA fragments were converted to sequencing library using the NEBNext® Small RNA Library Prep Set for Illumina® (NEB ref E7330S, USA) following the manufacturer’s recommendations. DNA library was quantified using a fluorometer (Qubit 3.0 fluorometer, Invitrogen, USA) and qualified using a High Sensitivity DNA chip on Agilent Bioanalyzer 2100. Libraries were multiplexed and subjected to high-throughput sequencing on an Illumina NextSeq2000 instrument with a 50 bp single-end read mode.

High quality raw sequencing reads (> Q30) were subjected to trimming using Trimmomatic v0.39 [83] with the following parameters: MINLEN:08, STRINGENCY:7, AVGQUAL:30, trimmed reads were used for alignment without further processing. Trimmed reads were aligned to the *S. pneumoniae D39* tRNA sequences extracted from the gtRNAdb [84] using bowtie2 v2.4.4 [85]in end-to-end mode (--no-unal --no-1mm-upfront-D 15-R 2-N 0 - L 10-i S,1,1.15 as other bowtie2 parameters), only uniquely mapped reads in positive orientation were retained for further analysis. 5’-reads’ extremities were counted for each RNA position in the reference; all further steps were performed in R/R-studio environment. After a-1 shift in the sequence position, since ligated 5’-P extremity is at the N+1 nucleotide in the RNA sequence, this reads’ count was used as the measure for intensity of cleavage at a given position. Four scores were used for analysis of AlkAnilineSeq raw data [80,81]: normalized cleavage (NCleavage), normalized count (NormCount), normalized G count (NormGcount) and stop ratio. Normalized cleavage corresponds to 1000x proportion of reads starting at a given position to the total number of reads mapped to a given RNA sequence. This score is less noisy but also less sensitive than others and is well-suited for detection of major cleavage events in RNA. In contrast, stop ratio (closely derived from ψ-ratio used for analysis of Ψ-seq data [86]) is calculated as a ratio of reads starting at a given position to the total number of reads passing (covering) it. This score is relatively sensitive, but also noisy. The last two scores used, NormCount and NormGcount use local normalization to the median of cleavage signals in ±5 nt window around of analysed position. NormCount uses all cleavage signals in the window, while signals corresponding to G residues are excluded from calculation of the normalization median for NormGcount. These scores represent the intensity of the cleavage at a given nucleotide compared to the local cleavage background in the adjacent RNA region. None of AlkAnilineSeq scores show linear dependence between the score’s value and stoichiometry of RNA modification, stop ratio has linear segment at very low modification levels (<5%), while the three other scores better represent the modification stoichiometry at higher modification levels in RNA (5-50%).

### Analysis of tRNA modifications by GLORI

GLORI is based on deamination and includes a glyoxal treatment that serves as a’protection’ step, selectively masking guanosine residues from nitrite (NO₂⁻)-induced deamination. This is followed by deprotection and reverse transcription to convert RNA into cDNA. The conditions used were as described in [87]. Briefly, 100 ng of tRNA samples were subjected to glyoxal treatment for 30 min at 50°C, followed by a boric acid treatment for 30 min at 50°C. The protected RNA is then subjected to nitrite-mediated deamination followed by RNA deprotection in a buffer containing 500 mM TEAA and formamide. RNA was then precipitated, end-repaired, and purified before being subjected to library preparation with NEBNext small RNA library following the manufacturer’s protocol. The quality and quantity of each library were assessed using a high-sensitivity DNA Chip on a Bioanalyzer 2100 and a Qubit 3.0 fluorometer. High!Zthroughput sequencing of the multiplexed libraries was performed on an Illumina NextSeq 2000 instrument in a 2×50 nt paired-end mode. Raw sequencing reads were inspected with FastQC and adapter sequence was removed by trimmomatic v0.39 [83]. Alignment to the A->G converted tRNA reference sequence (*S. pneumoniae D39* tRNA sequences) was done by Bowtie2. v2.4.4 [70] with slightly relaxed alignment stringency, allowing the retention of 1 nt-mismatched reads. Further analysis was done by *samtools mpileup* utility and counting the mismatch profile at every position in the reference. Since RNA A residues were de-aminated to inosines, sequencing reads are expected to contain only CGT nucleotides and thus align perfectly to A->G converted tRNA reference sequence. However, the residual A residues observed in the sequence data correspond to deamination-resistant modified As (m^6^A and all other N6-modified A, m^1^A was also detected). In the case of partial modification, both G and A are detected at a given position. Thus, the mismatch G in a reference to A in the sequencing read can be used as a score to measure a molar ratio of modified A in RNA (GtoA score). Raw AlkAnilineSeq and GLORI data are available at the European Nucleotide Archive under the accession number PRJEB96333.

## Results

### Combining LC-MS/MS, tRNAseq, and *in silico* analysis to predict tRNA modification genes in two model Streptococci

Over several years and studies [24,36], we performed LC-MS/MS analysis of bulk tRNA isolated from different *S. mutans* UA919 and *S. pneumoniae* D39 derivatives (13 independent *S. mutans* tRNA samples and 20 independent *S. pneumoniae* tRNA samples (**Table S1** and **Sup data 2BCDE**). By comparing retention times of chromatographic peaks in hydrolyzed tRNAs with those of synthetic standards, our analysis identified 31 modifications in *S. mutans* tRNAs and 32 modifications in *S. pneumoniae* tRNAs (**Supplemental Data 2BC** and **3**). Not all these modified nucleosides are from tRNAs. It is well known that modifications derived from contaminating 16S and 23S rRNA can be detected in high quantities in LC/MS analyses of nucleoside digests from tRNA preparations [20]. In addition, some modifications may originate from media contamination [85,86] or have non-enzymatic origins, such as m^1^A, m^3^C, or m^7^G [88]. Hence, LC/MS needs to be combined with tRNA-Seq and computational predictions to map the modifications on tRNA molecules and establish the links between tRNA modifications and their corresponding genes.

Two complementary strategies were employed to identify the tRNA modification enzymes encoded in the *S. mutans* UA159 genome. First, we used a curated set of 44 *B. subtilis* tRNA modification proteins (**Supplemental Data 1A**) to perform BLAST searches against the predicted proteomes of *S. mutans* UA159 (RefSeq Assembly: GCA_000007465.2) and *S. pneumoniae* D39 (RefSeq Assembly: GCA_000014365.1) to identify homologs. In parallel, we leveraged a curated list of orthologous groups from the COG and KEGG (KO) databases, mapped across ∼1,000 bacterial genomes [Reed C.J. and de Crécy-Lagard V. (unpublished) and **Supplemental Data 4**], to assess presence/absence patterns and cross-reference them with the BLAST hits. Integrating these datasets with our LC/MS profiles nucleosides in both *S. mutans* and *S. pneumoniae* (**Fig. 1**; **Supplemental Data 1BCD**).

We infer that several nucleosides detected by LC/MS—including acp^3^U, m²G, m²²G, m^3^C, m³U, mLJC, and mLJ,LJA, originate from rRNA or media contaminations, as no candidate tRNA-modifying enzymes were found for their incorporation. Putative rRNA-modifying enzymes could be identified for all of these, except for m^3^C and acp^3^U (**Supplemental Data 2F**). Traces of acp³U were observed exclusively in *S. pneumoniae*, likely due to contamination from the sheep blood in the TSAII-BA growth medium. Similarly, the traces of Q in *S. pneumoniae* samples are likely from media contamination (see section below). Both m^3^C and m^5^C were detected in both Streptococci, but no m^4^C was detected (**Supplemental Data 3**). The presence of SunL (16S rRNA m^5^C967-methyltransferase) homologs in both species (**Supplemental Data 2F)** suggests that m^5^C is indeed a rRNA modification. However, m^3^C had not been detected in bacterial RNAs to date [79], and homologs of RsmH that generally catalyze the insertion of m^4^C in rRNA were identified in both Streptococci (**Supplemental Data 2F**). Hence, in this case, the in silico and the LC/MS data did not match, warranting further investigations. As mLJCm is a conserved bacterial 30S rRNA modification [79] [and both Streptococcal genomes encode orthologs of *E. coli* RsmI (**Supplemental Data 2F**)—an enzyme responsible for 2’-O-methylation of C1402 in 16S rRNA [89]—we revisited our initial analysis and found that mLJCm was indeed present in our tRNA samples (**Supplemental Data 3).** To confirm this presence was due to contaminating rRNA fragments, we quantified mLJCm levels in bulk rRNA and tRNA extracted from *S. mutans* UA159 and indeed found them to be 750-fold higher in rRNA than tRNA (**Supplemental Data 2F and 3)**. The LC/MS analysis of nucleosides derived from *S. mutans* rRNA samples confirmed m²G, m²,²G, m³U, mLJC, and mLJ,LJA were only present in rRNAs, while Gm, Ψ, m^7^G, m^5^U, and possibly D were found in both rRNAs and tRNAs. This analysis also showed that m³C was only present in traces in rRNA and was certainly derived from a contamination or a non-enzymatic reaction in our tRNA samples. Finally, no Am was detected in rRNA; hence, we cannot rule out that this modification is enzymatically inserted in tRNA by TrmL or an as-yet-unknown enzyme.

Based on enzyme predictions and the LC/MS data, we estimate that 21 of the 31 nucleosides detected in *S. mutans* and 19 of the 32 in *S. pneumoniae* derive from tRNAs. AlkAnilineSeq and GLORI analyzes allowed us to specifically map the D, m^1^A, m^7^G, t^6^A, m^6^A and i^6^A modifications in *S. pneumoniae* tRNAs sequences (**Fig. 2C** and **Fig. S2**). Exact mapping of Ψ/Um/Cm/Gm and potentially Am in specific tRNAs would require additional analyses.

**Figure 2.**
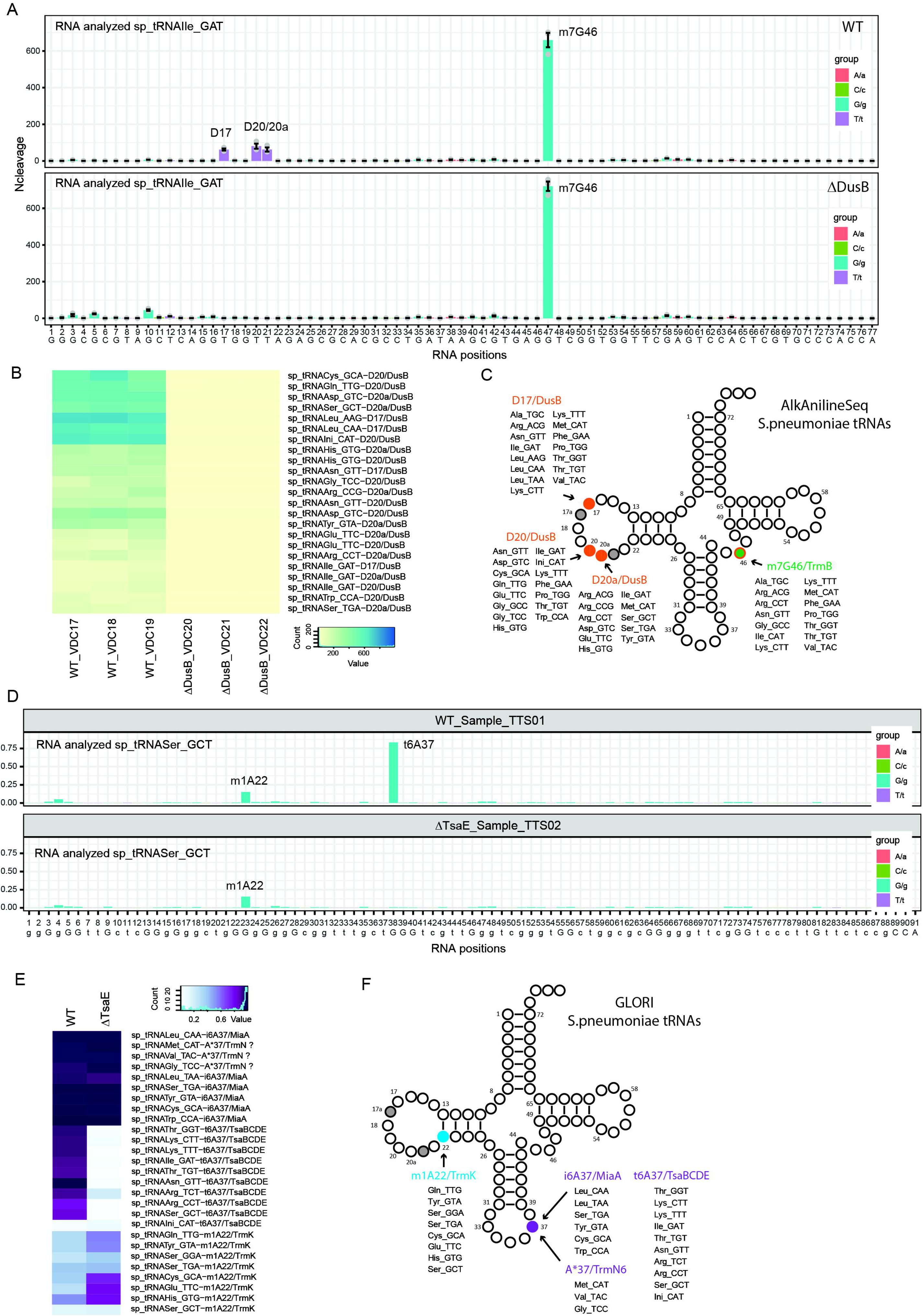
***S. pneumoniae* tRNA modifications mapped by sequencing-based modification detection methods.** (A) Alkaline Seq analysis of tRNA extracted from WT and Δ*dusB1 S. pneumoniae* Representative AlkAnilineSeq profiles showing signals for D (at positions 17, 20, 20a) and m^7^G (at position 46) in the WT and Δ*dusB1* strains. Ncleavage score is shown in the same scale, nucleotide color code and strain identity are shown on the right. AlkAniline-Seq signals are measures for biological triplicate (n=3) and the average value is shown, along with the error bar (calculated using the R ggplot2 package). Individual points for replicates are shown in grey. tRNA residues are numbered in sequential order, so position number may not correspond to conventional tRNA numbering (including 17a, 20a/b, and variable loop); (B) Heatmap for absolute Ncleavage values observed upon AlkAnilineSeq analysis of most modified D sites mapped in S. pneumoniae WT and *ΔdusB1* strains. The color code is shown at the bottom right, and the identity of the D site in tRNA is indicated. Data are shown for all replicates (n=3) for WT and ΔDusB strains; (C) Summary of all AlkAniline-Seq-detected modified residues in *S. pneumoniae* tRNAs. m1A residues are shown in orange, m^7^G are in green. Identity of tRNAs harboring these modifications is shown; (D) Representative GLORI profiles detecting m^1^A and i^6^A residues in *S. pneumoniae* tRNA. The signal in GLORI corresponds to non-deaminated (NO_2_-resistant) A residues (m^1^A and N6-substuituted A residues). Data for WT and *ΔtsaE* strain are shown. Nucleotide color code is shown on the right. tRNA residues are numbered in sequential order, so position number may not correspond to conventional tRNA numbering (including 17a, 20a/b, and variable loop); (E) Heatmap of the GLORI signals for all detectable modifications (m^1^A, m^6^A, i^6^A and t^6^A). The color code is shown on the top. Identity of tRNA modified site is indicated on the right. (F) Summary of all GLORI-detected modified residues in *S. pneumoniae* tRNAs. m^1^A residues are shown in light blue, N6-substituted A37 residues are in violet. A*37 indicates N6-substituted A which occurs in the sequence different from contexts AAA (for i^6^A) and UAA (for t^6^A), most likely these tRNAs harbor m^6^A modification. Identity of tRNAs harboring these modifications is shown.

The main difference between *B. subtilis* and *S. mutans* is the absence of the ms^2^i^6^A, ms^2^t^6^A, and ct^6^A-derived modifications and the corresponding enzymes (MiaB, MtaB, and TcdA) (**Fig. 1** and **Supplemental Data 1B**). In addition, the loss of one of the DusB2 orthologs and the absence of the alternate GTP cyclohydrolase IB/MtrA reduce the number of tRNA modification genes from 44 in *B. subtilis* to 39 in *S. mutans* (**Fig. 1** and **Supplemental Data 1B**). The same gene/modification losses were also observed in *S. pneumoniae,* but a further decay of complex modification pathways, such as the previously reported loss of MnmL [24] and the decay of the Q pathway described below, led to a final count of 33 tRNA modification genes in this respiratory pathogen (**Fig. 1** and **Sup data 1C**).

### DusB1 is responsible for all D synthesis at positions 17, 20, and 20a in *S. pneumoniae*

*S. mutans* and *S. pneumoniae* only encode a DusB1, whereas *B. subtilis* encodes DusB1 and DusB2 ([23] and **Fig, 1**). To determine which positions were modified by DusB1 in *S. pneumoniae,* the corresponding gene (*spd_2016*) was deleted, and AlkAnilineSeq was performed on tRNA extracted from the WT and Δ*dusB1* strains. We found that *dusB1* was responsible for the formation of D17, 20, and 20a, as all three modifications disappeared in the mutant (**Fig. 2AB and Fig. S2**). In the process, we mapped the D residues in 40 *S. pneumoniae* tRNAs **(Fig. S2).** We suggest that the DusB1 proteins of *S. mutans* and *S. pneumoniae* have similar catalytic profiles (**Fig. 1**), In *B. subtilis,* tRNA^Met^_CAU_ harbors a D at position 47 which was identified by LC/MS of the T1 fragments tRNA, that D47 can only be seen by AlkAnilineSeq in a *trmB* mutant, so that the m^7^G46 signal does not overshadow the D signal [23]. Hence, we cannot formally rule out the presence of D47 in *S. pneumoniae* in this tRNA. Recent analysis of *S. aureus* tRNA modification profiles [27] shows that D47 is not found in any tRNA; hence, we did not include this modification in our predictions (**Fig. 1**), even if the final call would require additional experiments.

During the mapping of D residues by AlkAnilineSeq, we also identified multiple m^7^G46 modifications in *S. pneumoniae* tRNAs (**Fig. 2AB and Fig. S2A** and, unexpectedly, several U8 also give an AlkAnilineSeq signal, presumably corresponding to the predicted s^4^U8. This observation was further confirmed by analysis of the mismatch score at those positions. As anticipated, all AlkAnilineSeq signals at position 8 were also detected as partial U->C transitions, which is a characteristic feature of s^4^U, base pairing with A, but also with G residues during the reverse transcription step (**Fig. S2B**). However, we believe that methods allowing specific s^4^U detection [90] should be used to map these modifications comprehensively and thus did not include them in our summary.

### *S. pneumoniae* salvages preQ_1_ and not preQ_0_, as predicted from metabolic reconstruction and stops at the intermediate oQ

Metabolic reconstruction of the *S. pneumoniae* Q synthesis pathway (**Fig. 3A and S3**, **Supplemental Data S1CD**) predicts that, unlike *S. mutans*, which is a Q prototroph, *S. pneumoniae* cannot synthesize preQ₀ de novo due to the absence of the *queCDE* genes [38]. However, the presence of *queT* and *queF* homologs suggests that *S. pneumoniae* can salvage preQ₀, reduce it to preQ₁, and incorporate it into tRNA via the signature enzyme TGT (**Fig. 3A**). Notably, homologs of *queG* and *queH*—two non-orthologous enzymes that catalyze the final step of Q biosynthesis—are missing. This indicates that the pathway in *S. pneumoniae* likely halts at the penultimate step: the formation of epoxyQ-tRNA (oQ-tRNA) from preQ_1_-tRNA catalyzed by QueA (**Fig. 3A and S3**). Such an incomplete Q pathway is highly unusual, with only one other documented case to date [91].

**Figure 3.**
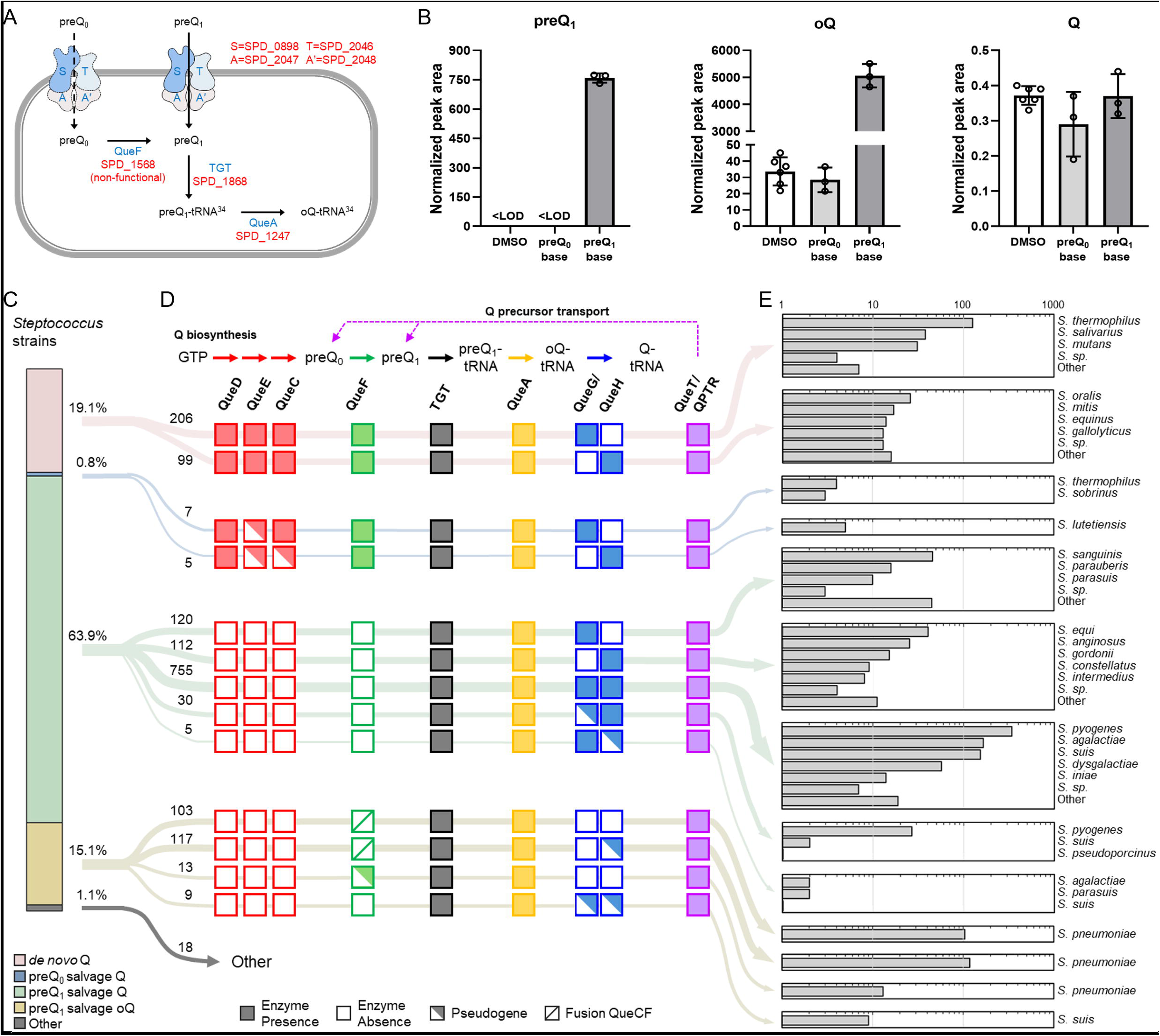
Decay of Q pathway in *S. pneumoniae* and other Streptococci. (A) Predicted Q pathway in *S. pneumoniae* D39, with protein names shown in blue and locus tags in red. The ECF transporters include 4 subunits: S, the substrate-specific transmembrane component (QueT); T, the energy-coupling module; A and A’, a pair of ATPase; (B) *S. pneumoniae* salvages preQ_1_ to synthesize oQ in tRNAs. The wild-type strain IU1824 was cultured in BHI medium supplemented with DMSO, 100 nM preQ_0_ base, or 100 nM preQ_1_ base. tRNAs were isolated, enzymatically digested, and analyzed by LC-MS/MS as described in the Materials and Methods section. For each analysis, 600 ng of hydrolyzed tRNAs was injected. The identities of the preQ₁ and Q peaks were confirmed by comparing them to the retention times of their synthetic standards. The identity of the oQ peak was confirmed through high-resolution LC-MS analysis, as detailed in the Materials and Methods and Results sections. Relative abundance for each modified ribonucleoside was calculated by normalizing its raw peak area to the sum of UV signals from the four canonical ribonucleosides. Reported averages and standard deviations are based on data from three independent growth experiments; (C) The distribution of different pathways of 7-deazaguanine modification in tRNA in 1,599 *Streptococcus*. The four major pathways are *de novo* Q biosynthesis (red), preQ_0_ salvage Q modification (blue), preQ_1_ salvage Q modification (green), and preQ_1_ salvage oQ modification (yellow), with the others (gray) consisting of Q gene patterns that were found in no more than two strains; (D) The Q gene distribution for each category. The Q biosynthesis and Q precursor transportation is depicted in the top panel. Each arrow represents one reaction catalyzed by the enzyme underneath. Dashed arrows represent Q precursor transmembrane salvage via transporter proteins. The presence and absence of each Q gene is indicated by the corresponding filled and open square. The presence of a pseudogene is indicated by a half-shaded square; (E) The composition (number of strains) of species for each type of Q gene pattern. For better visualization, less abundant species collapsed.

In our initial LC-MS analysis, *S. pneumoniae* was grown in BHI medium without added preQ₀ or preQ₁ (**Supplemental Data 2C**). Only trace amounts of Q were detected, likely due to media contamination. In contrast, oQ levels were ∼20-30 times higher in *S. pneumoniae* than in *S. mutans* (**Supplemental Data 2BC**). The identity of the oQ modification was confirmed using high-resolution LC-MS with targeted SIM and MS/MS analyses. SIM data showed excellent agreement between the observed m/z (426.1608) and the theoretical m/z (426.1619) for the protonated oQ ion, with a mass error < 3 ppm (**Fig. S4**). MS/MS analysis of m/z 426.1619 yielded major fragment ions at m/z 295.1035 (<1 ppm error) and 163.0616 (<2 ppm error), corresponding to the 7-methyl-7-deazaguanosine and 7-methyl-7-deazaguanine moieties of oQ, confirming its identity.

oQ levels in *S. pneumoniae* remained ∼10-fold lower than the Q levels observed in *S. mutans* (**Supplemental Data 2BC**). Because tRNAs were initially extracted from cells grown in undefined media with unknown concentrations of preQ₀/preQ₁, we repeated the experiment with 100 nM preQ₀ supplementation. However, oQ levels were unchanged (**Fig. 3B** and **Supplemental Data 2D**). This was unexpected given the presence of QueT (predicted to transport preQ_0_/preQ_1_) [36] and QueF predicted to reduce preQ_0_ into preQ_1_ [38]. Upon closer inspection, the *S. pneumoniae* D39 QueF was found to be lack catalytic residues: it is a fusion of an N-terminal QueC fragment and a C-terminal QueF domain lacking the catalytic cysteine essential for thiamide adduct formation with preQ₀ (**FigS5**) [92]. We then repeated the experiment with 100 nM preQ₁ instead. This led to a substantial (>100-fold) increase in oQ levels and high accumulation of preQ₁ in *S. pneumoniae* tRNA, while the trace Q levels remained unchanged (**Fig. 3B** and **Supplemental Data 2D**). The result was supported by the observation that all TGTs from *Streptococcus* have a substrate binding motif that resembles *E. coli* TGT which takes preQ_1_ as the substrate (**Fig. S6**).

In summary, unlike *S. mutans* UA159, which encodes a fully functional de novo Q synthesis pathway [36], *S. pneumoniae* D39 has lost the capacity to synthesize preQ_1_ but can still salvage this precursor to insert it into tRNAs. In this organism, the pathway also stops at the oQ-tRNA intermediate. These findings led us to conduct a comprehensive comparative genomic analysis of Q synthesis and salvage genes in sequenced Streptococci to evaluate the variability of the Q pathway across the entire clade.

### Plasticity of Q biosynthesis pathways in Streptococci as seen with numerous gene losses and pseudogene accumulation

We analyzed the distribution of queuosine (Q) synthesis and salvage genes in 1,599 *Streptococcus* genomes, selected based on genome completeness **(Fig. 3CD; Supplemental Data S1E**). 1598 strains are predicted to harbor a deazaguanine modification in tRNA, as they all encode a functional TGT enzyme, the remaining one is likely to be a sequencing error. QueA, the next enzyme in the pathway catalyzing the synthesis of oQ-tRNA from preQ_1_-tRNA [38] is also highly prevalent, making it unlikely that preQ_1_ is the final incorporated base, unlike in *Bartonella* species [93]. Fewer than 20% of the species analyzed synthesize Q *de novo*, a group that includes many *Streptococcus thermophilus* isolates, all examined strains of *Streptococcus salivarius*, *S. mutans*, *Streptococcus oralis*, and *Streptococcus mitis* (common inhabitants of oral cavity), and the group D streptococci *S. gallolyticus* and *S. equinus* which inhabit the GI tract of mammals (**Fig. 3CD**). Notably, *Streptococcus* species that synthesize preQ_1_ never encode both non-orthologous enzymes that can catalyze the hydrolytic reduction/deoxygenation of oQ (QueG or QueH). Instead, they encode only one or the other (**Fig. 3CD**). By contrast, 755 strains—mainly *Streptococcus pyogenes* and *Streptococcus suis*—encode both QueG and QueH, yet all of these are auxotrophic for preQ_1_. Most Streptococci (∼80%) must rely on preQ_1_ salvage (**Fig. 3C**). In line with the prevalence of preQ_1_ salvage, orthologs of QueT or QPTR are found in most Streptococci analyzed (**Fig. 3D**). Finally, we found that the termination of the pathway at the oQ step, which we experimentally validated in *S. pneumoniae* D39 (**Fig. 3B**), is predicted to occur in ∼15% of the strains analyzed. Many of these harbor *queH* and *queG* pseudogenes, detected in 131 and 39 strains, respectively (**Fig. 3CD**). Gene losses and decay were also prevalent with the *queF* genes. 116 unique QueF sequences of two types were found in 544 *Streptococcus* strains: canonical QueF and inactive QueCF fusion, like the one found in the *S pneumoniae* D39 strain (**Fig. 3CD** and **Fig. 4 and S5**). All the 220 analyzed *S. pneumoniae* genomes encode defective QueCF fusions (**Fig. 4B**)

**Figure 4.**
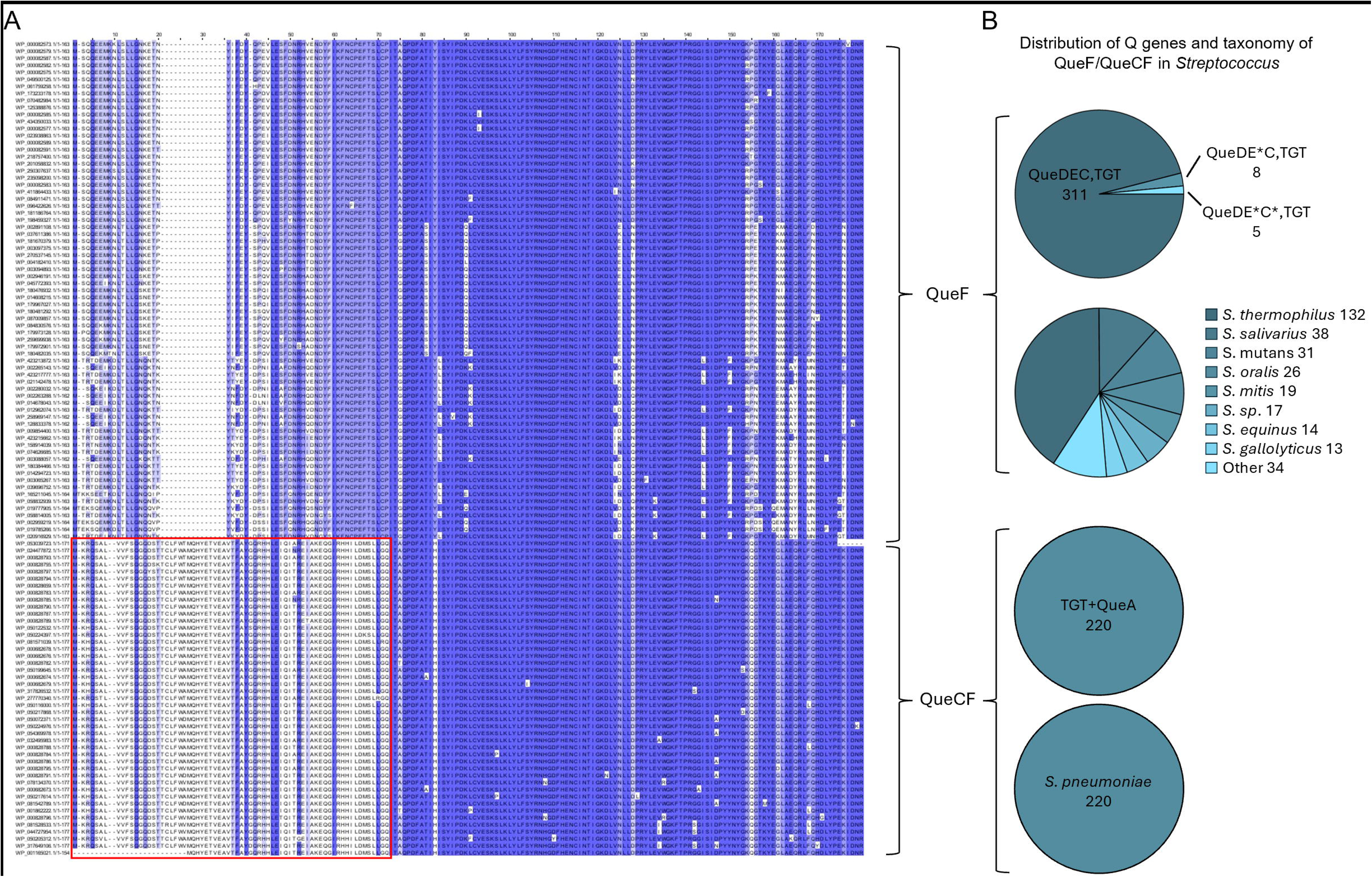
Two types of QueF proteins are found in Streptococcus. (A) The MSA of 116 unique QueF or QueCF from 544 Streptococcus strains. The boxed region indicates the N-terminus that do not align; (B) The corresponding Q genes and species of 544 QueF encoding Streptococcus strains. RefSeq accessions for all proteins listed are given in **Supplemental Data S1F.**

We also identified the Streptococci orthologs of QueK enzymes that hydrolyze the Q nucleoside to liberate the queuine (q) (**Supplemental Data 1E**), QueK proteins are members of the superfamily isine/uridine-preferring nucleoside hydrolase (IPR023186) family, but members of the QueK subgroup can be separated from the other members of this large family using Sequence Similarity Networks (SSNs) ([40] and **Fig. S7A**). Indeed, we found that QueK homologs from Streptococci are grouped with the experimentally validated QueK from *Clostridioides difficile* 630 (QueK Cd in **Fig. S7A**). Most genomes in this cluster lack *queDECF* genes but encode TGT. In addition, the majority of *queK* genes in this cluster, including those from *Streptococcus*, are next to genes encoding QueT family transporters (**Fig. S7B**). Hence, it is likely that QueK orthologs in *Streptococcus* function as queuosine hydrolase. However, no QueL orthologs could be identified in the analyzed *Streptococcus* genomes (**Supplemental Data 1E**). QueA orthologs are always present (Fig. 3CD), and the encoded Tgt proteins are all of the canonical preQ_1_ inserting/bacterial type [94] (Fig. S6). Hence, it is not clear what the fate of the released q base would be in Streptococci. One cannot rule out the possibility that a non-orthologous enzyme yet to be discovered replaces QueL to generate preQ_1_ in these organisms.

To investigate the potential impact of oQ on synonymous codon usage of the GUN Q-dependent codons (Tyr, His, Asp and Asn), we compared codon usage of 26 of the 234 *S. pneumoniae* strains in our dataset predicted to harbor oQ each to a closely related Streptococci (>92% identity) not belonging to the *S. pneumoniae* clade (**Supplemental Data 1F**). We found that *S. pneumoniae* strains exhibited a higher NAU-to-NAC ratio (**Fig. S8A**), which was primarily attributed to the Tyr and His codon biases (**Fig. S8C-F**). However, this difference could be a result of difference in GC content (Fig. S8B) (**Supplemental Data 1F**) as this can be a driver the choice of the wobble base [95]and more extensive analyses will be required to separate these effects.

### Survey of the essentiality of tRNA modification genes in *S. mutans* and *S. pneumoniae* reveals differences between the two clades

Based on published studies [96–100] and TnSeq data generated as described in the methods section, we compiled essentiality data for all predicted *S. mutans* UA150 and *S. pneumoniae* D39 tRNA modification genes (**Supplemental Data 1BC**). As expected from the literature [17], *trmD* and *tilS* were essential in both species. *IscS1* and *mnmA,* both involved in the synthesis of s^2^U34 were also essential in both species, whereas they can be deleted in both *E. coli* and *B. subtilis* [101,102]. Discrepancies between TnSeq data, which often report *mnmA* as essential, including in *E. coli* [103], and targeted deletion studies showing that *mnmA* is frequently dispensable [104] can be explained by the severely impaired growth of *mnmA* mutants [101,102] which leads to their counterselection in pooled populations. It appears that *mnmA* is essential in Streptococci, as directly targeting the genes for deletion in *S. mutans* was unsuccessful (**Supplemental Data 1B).**

The *tadA* gene is essential in *E. coli* but not in organisms such as *B. subtilis* [105,106] or *S. aureus* [Marzi], even if it is not totally clear why the unmodified tRNA^Arg^_ACG_ can decode all CGN codons in *B. subtilis* and not in *E. coli* [105]. TadA is not essential in *S. mutans,* and the two current studies utilizing two different growth media disagree on the essentiality of the gene in *S. pneumoniae* (**Supplemental Data 1BC**); therefore, additional targeted genetic experiments will be necessary.

If no gene was found to be essential only in *S. mutans*, several genes were essential only in *S. pneumoniae* including *mnmEG* and *tsaBCDE.* MmnEG encoding genes are generally not essential in Streptococci, even if their absence leads to virulence defects [31–34]. We had previously confirmed that t^6^A was not essential in *S. mutans* as *tsaB*, *tsaC* and *tsaE* deletions were viable and devoid of t^6^A [37]. In contrast, all four t^6^A pathway-related genes seem to be essential in *S. pneumoniae* (**Supplemental Data S1C, Fig 5A and S9**).

**Figure 5.**
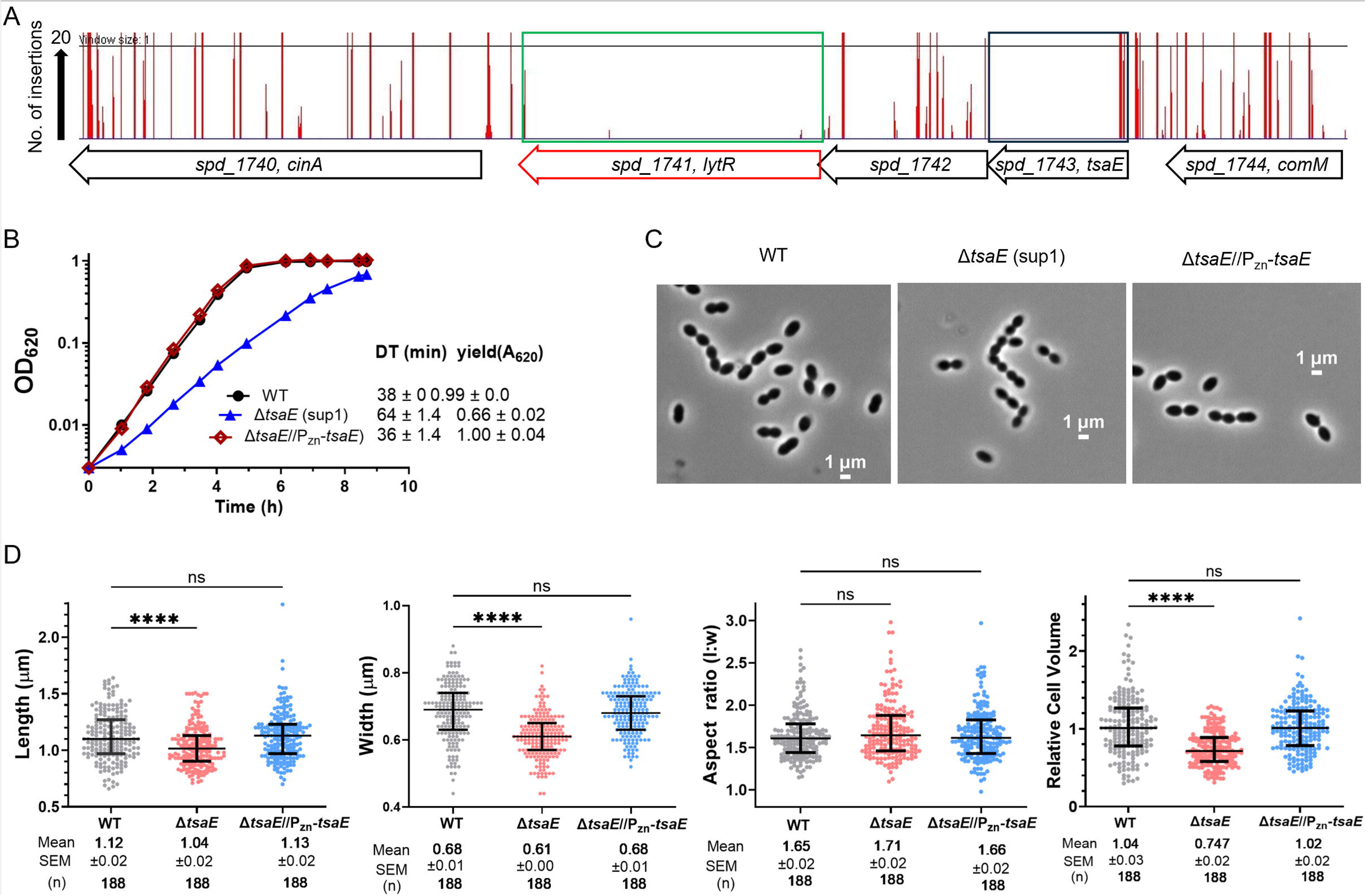
TsaE is essential in *S.pneumoniae* D39. (A) Mini-Mariner *Malgellan6* Tn-Seq transposon insertion profile for the genome region covering *spd_1740* (*cinA*), *spd_1741* (*lytR*), *spd1742*, *spd_1743* (*tsaE*) and *spd_1744* (*comM*) in the genomes of the unencapsulated WT parent (D39 Δ*cps rpsL1*, IU1824) strain growing exponentially in BHI broth in 5% CO_2_. The same WT Tn-seq insertion profile was obtained for encapsulated D39 strain IU1781 grown in BHI broth (data not shown). *In vitro* transposition reactions containing purified genomic DNA, *Magellan6* plasmid DNA, and purified MarC9 mariner transposase, transformation, harvesting of transposon-inserted mutants, growth of pooled insertion libraries exponentially in BHI broth, NextSeq 75 high-output sequencing, and analysis were performed as described in *Materials and Methods* based on [73]. Tn-insertions were recovered in the regions encoding *spd_1740*, *spd_1742* and *spd_1744*, but not in *spd_1741*(*lytR*) and *spd_1743* (*tsaE*). (B) Representative growth curves of the WT parent (IU1824), Δ*tsaE (*sup1) (IU18824), and Δ*tsaE* complementation strain (IU18963, Δ*tsaE* markerless//Δ*bgaA::*P*_zn_-tsaE*). *tsaE* deletion strains containing suppressor mutation *asnS*(D121A) are viable but have higher doubling time and lower yield in BHI growth media, and the growth phenotypes are complemented by ectopic *tsaE* expression. IU18824 was whole-genome-sequenced to contain *asnS*(D121A) mutation. IU1824 and IU18824 were grown overnight in BHI broth with no additional Zn^2+^/Mn^2+^ addition. IU18963 was grown overnight and day of experiment in BHI broth with additional 0.1 mM (Zn^2+^/(1/10) Mn^2+^) to induce expression of *tsaE* from the ectopic *bgaA* site. Overnight cultures were diluted to OD_620_ ≈0.003 in the morning in BHI broth for IU1824 and IU18824 and in BHI broth containing 0.1 mM Zn^2+^/(1/10)Mn^2+^ for IU18963. Doubling times (DT) and maximal growth yields (OD_620_) (averages ± SE) were obtained during 9 h of growth from 2 independent experiments. (C) and (D) Δ*tsaE* cells with an *asnS* mutations are smaller but with the same shape as WT strain, and the Δ*tsaE* cell morphology phenotype is complemented by ectopic expression of *tsaE*^+^ (C) Representative phase-contrast images taken at mid-log growth between 3 to 3.5 h for IU1824 and IU18963 (OD_620_ between 0.1 to 0.2), and between 3.5 to 4.0 h for IU18824 (OD_620_ between 0.03 to 0.05). Scale bar = 1 μm. (D) Scatter plots of cell lengths, widths, aspect ratios (cell length to width) and relative cell volumes of the above strains. Lines represent median and 25 and 75 percentiles. IU18963 was grown with 0.1 mM Zn^2+^ (Zn^2+^/(1/10) Mn^2+^) shown in (B). Pvalues were obtained by one-way ANOVA analysis (GraphPad Prism, Kruskal-Wallis test). **** and ns denote p<0.0001, and not significant, respectively when compared to WT.

### t^6^A synthesis genes are essential in *S. pneumoniae*, but a suppressor mutation in asnS make *tsaE* deletion strains viable

We confirmed the essentiality of the *tsaE* gene in *S. pneumoniae* by attempting to transform Δ*tsaE* amplicons into the unencapsulated IU1824 (D39 Δ*cps rpsL1*) or IU1945 (D39 Δ*cps rpsL^+^*) background. No transformants were recovered in the wild-type strains, whereas normal colony formation was observed in a strain expressing *tsaE* ectopically under the control of a zinc-inducible promoter (**Table 1**). WT strains transformed with *tsaE* deletion amplicons did appear after 40 hours of growth, most certainly having acquired suppressor mutations. Growth of Δ*tsaE* clones was affected compared to WT, and the growth defect was complemented by *tsaE* expression in trans (**Fig. 5B and Fig. 6AB**). The Δ*tsaE* cells were slightly smaller but the same shape as WT cells (**Fig. 6CD**). Whole genome sequencing of four independent Δ*tsaE* strains obtained from two different genetic backgrounds and two deletion markers revealed that all of them had acquired a mutation [*asnS*(D121A)] encoding asparaginyl-tRNA synthetase (AsnRS) (**Table 2**). To confirm that the presence of *asnS*(D121A) mutation can suppress the lethality of Δ*tsaE,* we constructed an isogenic set of strains expressing *asnS^+^* in the native chromosomal locus and an ectopic copy of either *asnS*^+^ or *asnS*(D121A). The presence of an ectopic copy of *asnS*(D121A), but not WT *asnS* allows the growth of a Δ*tsaE* strain **(Table 2 and Fig. 6B)**. This result indicates that *asnS*(D121A) is a gain-of-function mutation, consistent with the structural model described below (**Fig. 7**). Analysis of t^6^A levels in these strains by LC-MS (**Fig. 6CD**) and tRNA seq (GLORI) (**Fig. 2DE**) showed that t^6^A was indeed absent in the Δ*tsaE* strains.

**Figure 6.**
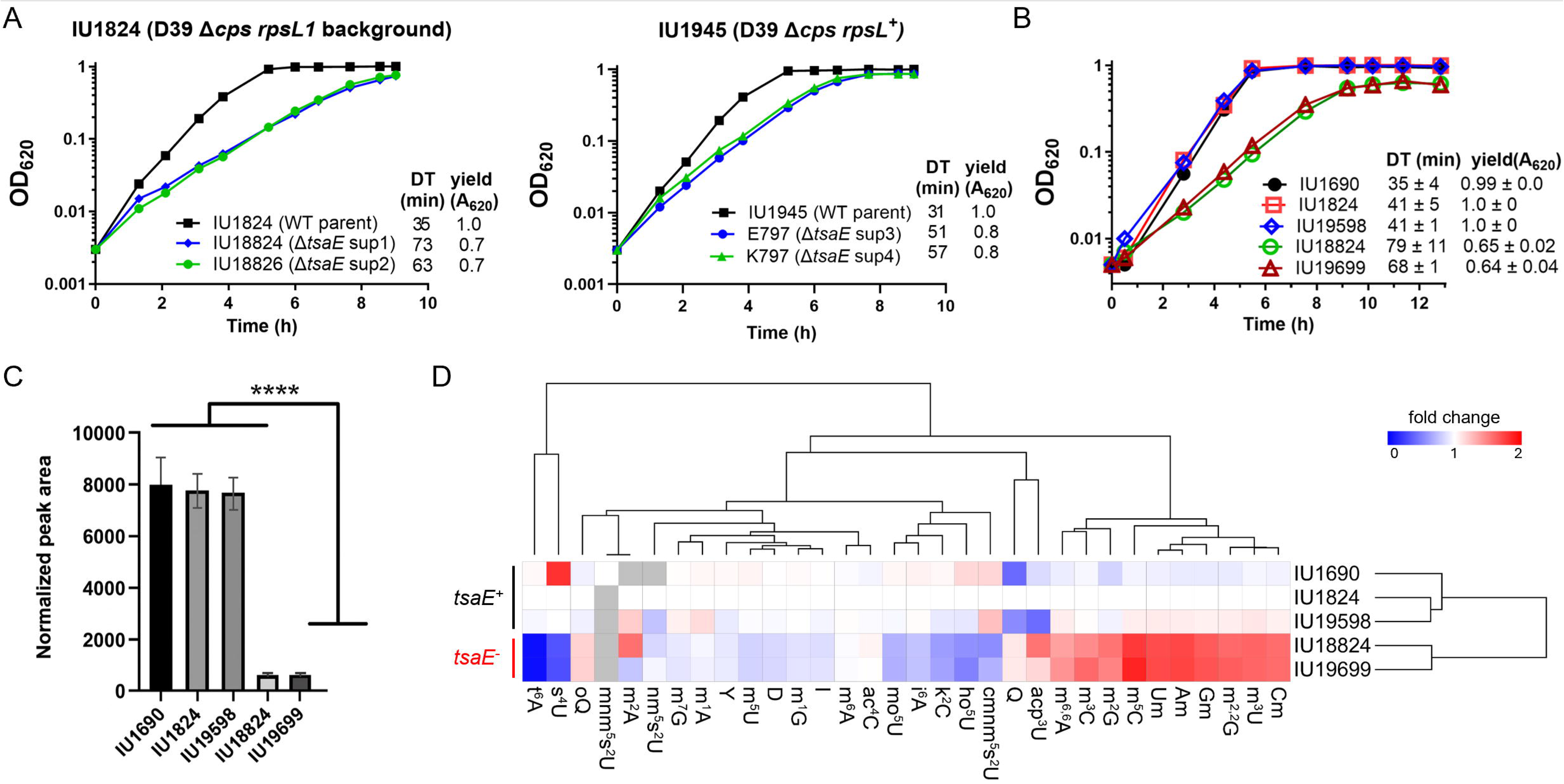
Suppressor mutations in *asnS* gene allow growth of *S. pneumoniae* D39 t^6^A-deficent strain. (A) *tsaA* deletion strains containing suppressor mutation *asnS*(D121A) are viable but have longer doubling time and lower yield in BHI growth media. Left panel, representative growth curves of the WT parent (IU1824, D39W *rpsL1* Δ*cps*) and mutants derived from this background, Δ*tsaE (*sup1) (IU18824, IU1824 Δ*tsaE::*P*_c_-erm* sup1), and Δ*tsaE (*sup2) (IU18826, IU1824 Δ*tsaE::*P*_c_-kan-rpsL* sup2). Right panel, representative growth curves of the WT parent (IU1945, D39W *rpsL^+^* Δ*cps*) and mutants derived from this background Δ*tsaE (*sup3), (E797, IU1945 Δ*tsaE::*P*_c_-erm* sup3), and Δ*tsaE (*sup4) (K797, IU1945 Δ*tsaE::*P*_c_-kan-rpsL* sup4). IU18824, IU18826, E797 and K797 were whole-genome-sequenced to contain *asnS*(D121A) mutation. Overnight cultures grown in BHI media were diluted to OD_620_ ≈0.003 in the morning in BHI broth. Doubling times (DT) and maximal growth yields (OD_620_) (averages ± SE) were obtained during 9h of growth from one experiment. (B) Representative growth curves of IU1690, IU1824, IU19598, IU18824, and IU19699. Expression of *asnS*(D121A) in the native genetic locus or at an ectopic site allows growth of *tsaE* deletion strains. IU1690 is D39W *cps*^+^ reference strain. IU1824 is D39 Δ*cps rpsL1*, the WT parent strain for IU19598, IU18824, and IU19699. IU19598 is IU1824 Δ*bgaA*::P-*asnS* (D121A), harboring ectopic expression of *asnS*(D121A). IU18824 is IU1824 Δ*tsaE*::P_c_-*erm* sup 1 (with suppressor mutation *asnS* [D121A]). IU19699 is IU1824 Δ*bgaA*::P-*asnS* (D121A) Δ*tsaE::*P_c_-*erm.* This strain is reconstructed to confirm *asnS* (D121) suppression of Δ*tsaE*::P_c_-*erm*. (C) Relative abundance of t^6^A in tRNAs isolated from IU1690, IU1824, IU19598, IU18824, and IU19699. tRNAs were isolated, enzymatically digested, and analyzed by LC-MS/MS as described in Methods section. For each analysis, 600 ng of hydrolyzed tRNAs was injected. The identity of the t^6^A peak was confirmed by comparing to the retention time of its synthetic standard. Relative abundance of t^6^A was calculated by normalizing its raw peak area to the sum of UV signals from the four canonical ribonucleosides. Reported averages and standard deviations are based on data from two independent growth experiments. Differences in t^6^A levels among the five strains were assessed using a one-way analysis of variance (ANOVA) at a significance level of *p* < 0.05, followed by Tukey’s post-hoc test for multiple comparisons. Statistically significant differences were indicated by \*\*\*\**p* < 0.0001. Statistical analysis was performed using GraphPad Prism 9. (D) Changes in relative abundance of modified ribonucleosides in tRNAs isolated from IU1690, IU1824, IU19598, IU18824, and IU19699. Each grid cell represents the mean fold-change for each modified ribonucleoside in IU1690, IU19598, IU18824, and IU19699 relative to IU1824, calculated from two biological replicates. An exception applies to mnm^5^s^2^U, for which fold-changes were calculated relative to IU1690. Grey cells indicate LC-MS/MS signals below limit of detection. Hierarchical clustering analysis was performed using Morpheus (https://software.broadinstitute.org/morpheus/), employing the Euclidean distance metric and the average linkage method to group both strains and modified ribonucleosides.

**Figure 7.**
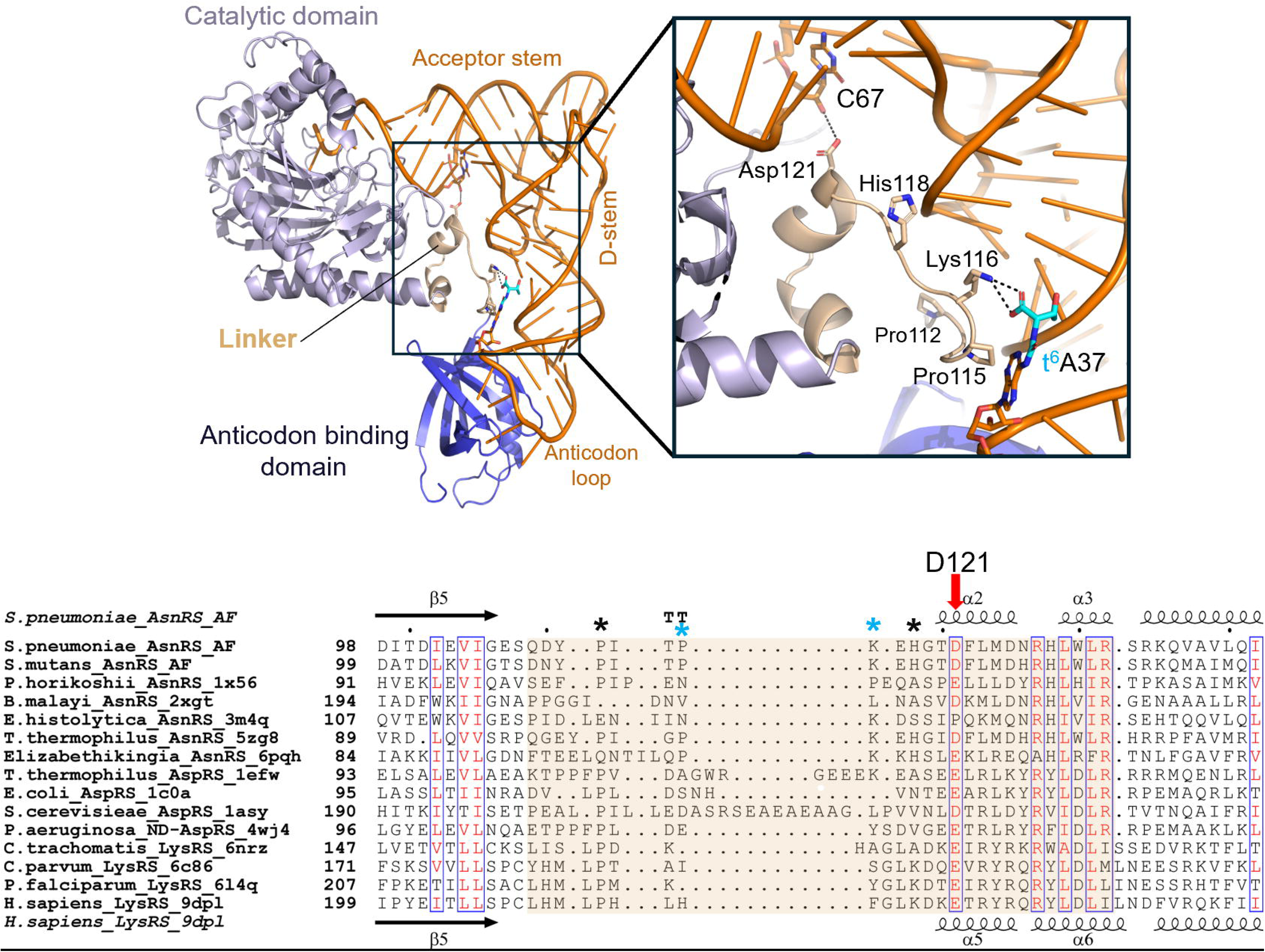
**Structural model of AsnRS/tRNA^Asn^ complex. (**A) model of t^6^A-modified tRNA^Asn^ bound to *Streptococcus pneumoniae* asparaginyl-tRNA synthetase (*Sp*AsnRS, purple), highlighting the inter-domain linker region (tan), potential interactions of linker residues with C67 in the base of the tRNA acceptor stem and with t^6^A37. The threonylcarbamoyl moiety of t^6^A37 is highlighted in cyan. (B) Structure-based multi-sequence alignment of AsnRS, ND-AspRS, and LysRS in the linker region (tan highlight). The conserved Asp/Glu corresponding to Asp121 of *Sp*AsnRS is indicated with a red arrow. Residues putatively involved in D-stem and acceptor arm are indicated with black and blue stars, respectively. PDB IDs for the structures used in generating the alignment are indicated in the sequence headers. AF: AlphaFold model.

**Table 2.**
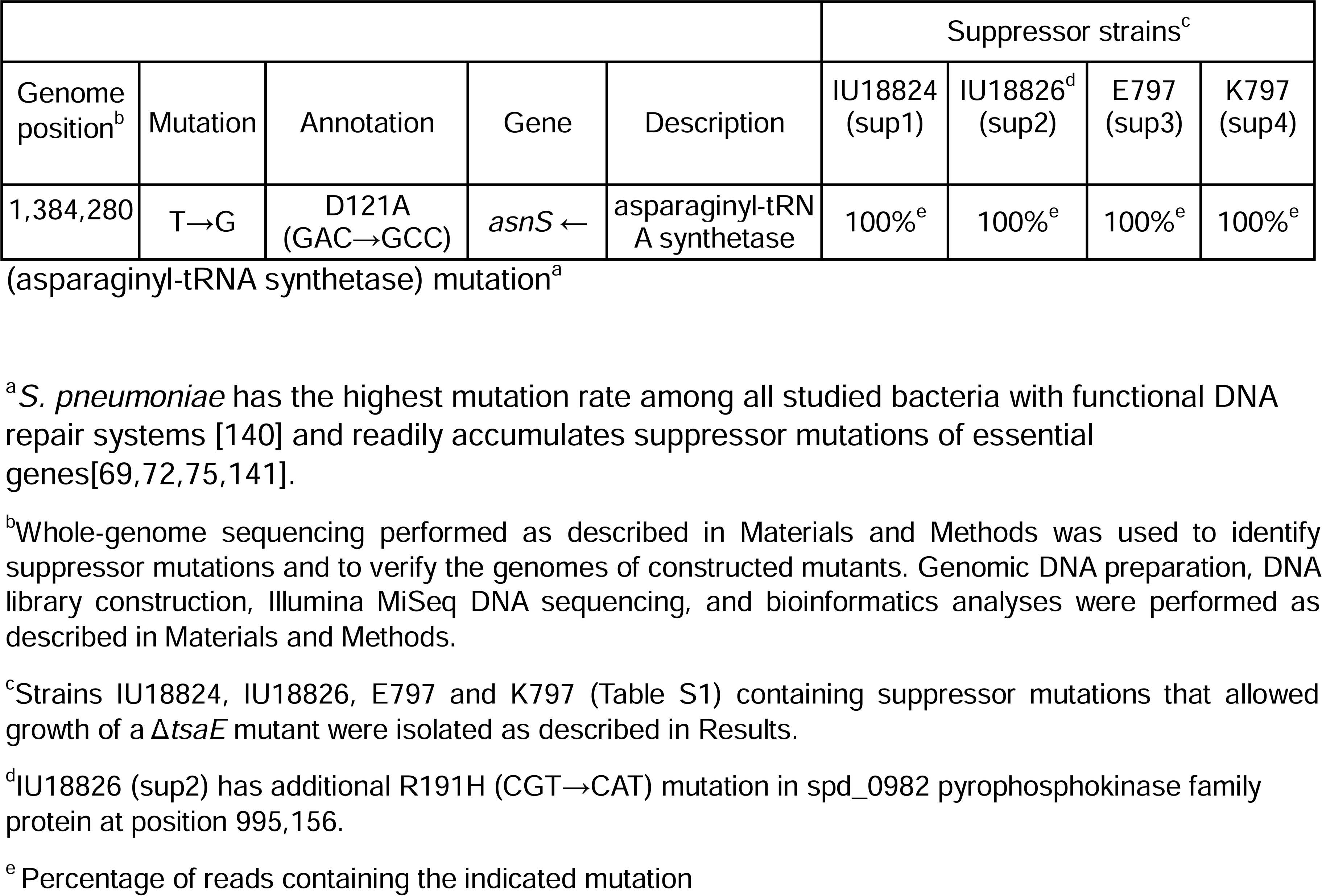
*tsaE* suppressor mutants of *S. pneumoniae* D39 contain *asnS*(D121A)

Profiling of the other rRNA/tRNA modifications by LC/MS showed that relative levels of all predicted rRNA modifications increased by 50% while all tRNA modifications (other than t^6^A) decreased by 20% (**Supplemental data 2C** and **Fig. 6D**). Based on well-established mechanistic links between cell stress and rRNA degradation [107,108], we believe the growth rate defect of the *tsaE* mutant strains leads to rRNA degradation and contamination of the tRNA preparations, which explains the changes in modification profiles.

### Structural modeling suggests a role for t^6^A37 in tRNA/AsnRS interactions

To gain structural insight into the role of Asp121 in *S. pneumoniae* AsnRS (*Sp*AsnRS) function, we constructed a model of its complex with t^6^A-modified tRNA^Asn^ (**Fig. 7**). Although crystal structures of AsnRS from several organisms are available, none include bound tRNA. The only available experimental structure of tRNA^Asn^ is that of *Pseudomonas aeruginosa* tRNA^Asn^ (*Pa*tRNA^Asn^) in complex with non-discriminating AspRS (*Pa*ND-AspRS, PDB ID 4WJ4 [109]), but this tRNA lacks the t^6^A37 modification. By contrast, recent high-resolution cryo-EM structures of human LysRS bound to ms^2^t^6^A37-modified human tRNA^Lys3^ (PDB IDs 9DPL and 9DPA [110]) provide insight into how a class IIb synthetase accommodates t^6^A37 in the anticodon loop and remodels the loop for recognition.

We used the AlphaFold2 model of *Sp*AsnRS (*Sp*AsnRS) [111], which closely resembles AsnRS crystal structures from diverse organisms, including *Thermus thermophilus* (PDB ID 5ZG8), *Elizabethkingia sp.* (PDB ID 6PQH, *Pyrococcus horikoshii* (PDB ID 1X54 [112]), *Brugia malayi* (PDB ID 2XGT [113]), *Entamoeba histolytica* (PDB ID 3M4Q) and *Homo sapiens* (PDB ID 8H53 [114]). The *Sp*AsnRS model superposes with these orthologs with r.m.s.d. values of 0.945-1.564 Å over 294-333 C_α_ atoms and aligns well with other class IIb synthetases including

*E. coli* AspRS (r.m.s.d. 2.23 Å over 269 C_α_ atoms) and *E. coli* LysRS (r.m.s.d 1.5 Å over 244 C_α_ atoms). *Pa*tRNA^Asn^ was positioned onto *Sp*AsnRS by superposing *Pa*ND-AspRS bound to tRNA^Asn^ with *Sp*AsnRS (r.m.s.d. 1.8 Å over 261 C_α_ atoms, 25% sequence identity, **Fig. S10A**). In parallel, the cryo-EM structure of human LysRS/tRNA^Lys3^ was superposed onto *Sp*AsnRS to inform the placement and orientation of the anticodon-binding domain relative to a t^6^A37-modified anticodon loop (r.m.s.d 5.3 Å over 311 C_α_ atoms, 18% sequence identity, **Fig. S10B**). Together, these superpositions yielded a composite model of the *Sp*AsnRS/tRNA^Asn^ that integrates the conserved fold of class IIb synthetases with structural evidence for anticodon remodeling and t^6^A37 accommodation (**Fig. 7A**).

AsnRS, AspRS and LysRS-II are all class IIb synthetases that act on ANN-decoding tRNAs, substrates of the t^6^A pathway. They share a conserved architecture comprising an N-terminal anticodon binding domain, a C-terminal catalytic domain, and an intervening 25-30 residue linker/hinge region formed by a flexible, proline-rich loop and two short α-helices. This linker articulates the catalytic and anticodon-binding domains, making direct tRNA contacts and positioning the acceptor end in the active site through interactions with the acceptor arm and the D-stem minor groove [110,115–117]. Our model suggests that these interactions are also possible in *Sp*AsnRS/tRNA^Asn^. For example, Pro112 and His118 in *Sp*AsnRS mimic the D-stem contacts made by Pro214 and Lys221 in human LysRS, respectively (**Fig. S10B**). In addition, Asp121 in *Sp*AsnRS aligns with Glu224 in human LysRS and Glu122 in *Pa*ND-AspRS, each observed to form a hydrogen bond with the 2′-OH of ribose 67 at the base of the acceptor stem, a contact that is largely conserved among class IIb synthetases, consistent with the conservation of this acidic residue across the family (**Fig. 7B**).

Linker residues in LysRS engage the modified base ms^2^t^6^A37 in the anticodon loop: Phe218 stacks against, and His217 hydrogen bonds with, the threonylcarbamoyl group of ms^2^t^6^A37 (Fig. S10B). Our model places *Sp*AsnRS residues Lys116 and Pro115 in analogous positions, suggesting similar interactions with t^6^A37 in t^6^A-modified tRNA^Asn^. When A37 is unmodified, this anticodon-loop contact is lost. Combined with the Asp121Ala substitution, which disrupts the acceptor-stem contact, the linker may be released to adopt an alternative conformation that maintains D-stem interactions. This flexibility provides a plausible route for *Sp*AsnRS to function with unmodified tRNA^Asn^.

Based on the above structural analysis and the suppression phenotypes of strains carrying *asnS*(D121A) mutation, we hypothesize the following suppressor mechanism. t^6^A at position 37 of tRNA acts as a strong positive determinant with WT SpAsnRS, similar to the case of t^6^A and *E. coli* IleRS [43]. WT SpAsnRS can form a productive complex with t^6^A-modified tRNA, but not with unmodified tRNA. Since t^6^A-modified tRNA^Asn^ is absent in a *tsaE* deletion mutant, the lack of Asn-charging leads to lethality of Δ*tsaE* cells. We hypothesize that *Sp*AsnRS(D121A) encoded by the suppressor mutation *asnS*(D121A) allows the interaction of the synthetase with unmodified tRNA, leading to Asn charging for translation. However, this reaction is less efficient than that carried out by WT AsnRS with modified tRNA, resulting in slower growth in the suppressed strain. This model is consistent with the result showing *asnS*(D121A) as a gain-of-function mutation. The presence of AsnRS with this amino acid change is sufficient to allow cell growth, independent of the presence or absence of WT AsnRS.

## Discussion

It is also well established that pathogenic bacteria tend to streamline their genomes compared to their environmental ancestors [118]. Accordingly, *S. mutans* and *S. pneumoniae* retain 87% (39/45) and 78% (35/45), respectively, of the tRNA modification genes found in *B. subtilis (***Fig. 1** and **Supplemental Data 1ABC**). What was unexpected, however, was that the majority of the lost genes encode Fe-S cluster –requiring enzymes. Specifically, 80% of the missing genes in *S. mutans* and 60% in *S. pneumoniae* fall into this category, including *mnmL*, *queG*, *queE*, *mtaB*, *miaB*, and *tcdA* (**Fig. 1** and **Supplemental data 1ABC**). Previous studies had already noted that many widespread Fe–S-dependent metabolic proteins are absent in *S. mutans* and in other Gram-positive species [119]. In addition, the SUF system responsible for Fe–S cluster assembly is not essential in *S. mutans*, unlike in *B. subtilis*, because the essential Fe–S enzymes of isoprenoid synthesis (IspH and IspG) have been replaced in *S. mutans* by the Fe-S independent mevalonate pathway [120]. Hence, we see that the selective pressure in both these organisms has led to shedding hyper-modifications that require Fe–S enzymes, while still maintaining modifications in the same positions. *S. mutans* is exposed to a variety of reactive oxygen species (ROS) in the oral cavity, such as H_2_O_2_ production by niche-competing early colonizers of dental plaque biofilm (*S. gordonii*, *S. sanguinus*) [121,122], periodontal inflammation [123], hypothiocyanite production by host salivary lactoperoxidase [124], and use of H_2_O_2_-containing dental care products. Given that Fe-S clusters are an important source of hydroxyl radical generation via Fenton chemistry during exposure to oxidative stress, it makes sense for *S. mutans* to minimize its reliance on Fe-S containing enzymes required for central physiological functions such as tRNA modification. One such specific example is the Q synthesis pathway. *S. pneumoniae* has lost the two Fe-S dependent enzymes of this pathway: QueE [125], involved in the preQ_0_ precursor synthesis, and QueG, involved in the last step, the reduction of oQ. However, we show that the pathway has not been lost and oQ accumulates in *S. pneumoniae* tRNA. Even if the Q modification has been repeatedly and independently lost in many clades [38], all the circa 1599 Streptococci genomes analyzed are predicted to still carry Q, or the Q precursor oQ, in target tRNAs, reinforcing its critical role in translation accuracy in this clade. This comparative work emphasizes the difficulty in predicting active pathways through metabolic reconstruction. Indeed, as we show here, pseudogenes can create false positive hits that can skew the pathway predictions: in this case, we predicted preQ_0_ salvage in *S. pneumoniae* when in reality, preQ_0_ is salvaged in this organism because the QueF homolog is not functional (**Fig. 3**).

The translation machinery is a classical target for antibacterial compounds with many known antibiotics targeting one aspect of translation or another (such as aminoacyl-tRNA synthetases, ribosome, translation factors)[126]. One aspect of translation that has not been targeted is tRNA modification, for two main reasons. First, it is only in the last decade that many of the genes/enzymes involved in tRNA modification have been identified at least in one model organism from each kingdom [6]. Second, only a handful of modification genes are essential in most pathogenic bacteria. These are the genes that introduce m^1^G37, k^2^C34, and t^6^A37. TrmD, the m^1^G37 methylase, was the first to be identified as an antibacterial target because it is very different from its human counterpart (Trm5), and inhibitors have already been identified [18,127]. TilS, the enzyme that introduces the k^2^C modification, is an obvious target as it is absent in eukaryotes [128]. TilS and TrmD are both essential in both *S. mutans* and *S. pneumoniae* (**Supplemental data S1BC**), hence any antibiotic targeting these enzymes should inhibit the growth of both these pathogens. The situation is not as clear-cut for t^6^A synthesis.

The differences between the bacterial and eukaryotic t^6^A synthesis machinery open the possibility of targeting this pathway for antibacterial compounds [41,42]. While the first enzyme of the pathway that produces threonylcarbamoyl adenylate, (TsaC in *E. coli*, Tsc2/Sua5 in yeast, YRDC in human) is very conserved across kingdoms [76], the second step of the pathway, the transfer of the threonylcarbamoyl moiety to the target adenosine, varies greatly (see [43] for review), with only one protein family (TsaD/Kae1/Qri7) in common between the different pathways. Mitochondria just require Qri7 to complete t^6^A synthesis, whereas Bacteria require TsaB and TsaE in addition to TsaD, and the whole KEOPS complex catalyzes this reaction in all eukaryotes [41,42]. Because of their prokaryotic-specific essentiality, the *tsaBE* genes had been identified as potential antibacterial targets before their role in t^6^A synthesis was even established [129–134] and inhibitors of TsaE were developed based on its ATP-binding capabilities [135]. TsaE is essential in many bacterial pathogens, including nearly all ESKAPE pathogens (**Supplemental Data S1J**); however, it can be deleted in some pathogenic bacteria, such as *S. mutans*. We showed here that *tsaE* is essential in *S. pneumoniae* but a suppressor mutation in the *asnS* gene encoding AsnRS was readily selected, suggesting a role for t^6^A in AsnRS function in this organism. The mutation results in a D121A substitution in the coding sequence, which could increase flexibility of a linker region and compensate for the loss of anticodon loop interactions that normally depend on t^6^A37. In contrast, *S. mutans* AsnRS (*Sm*AsnRS) retains the wild-type sequence in a *ΔtsaE* strain lacking t^6^A, indicating that t^6^A37 is not critical for *Sm*AsnRS function. D121 is conserved in *Sm*AsnRS, and the protein shares 80% sequence identity with *Sp*AsnRS, including in the linker region (**Fig. 7B**), with predicted 3D structures that are virtually identical. Notably, *S. pneumoniae* and *S. mutans* tRNA^Asn^ molecules differ by a single base pair (C5:G69 versus G5:C69, respectively; **Fig. S10C**), located adjacent to C67 in our docking model and near the contact point for D121. This difference may influence local geometry at the base of the acceptor stem in ways not evident in current structures, but the structural basis for the different requirements for t^6^A37 remains unclear. It is worth noting that in our study only one unique *asnS* mutation within the linker region was obtained among four independent mutants. In comparison, evolution repair studies of *E. coli* growth of *trmD* mutant strains show that growth of *trmD* mutants can be largely restored by single mutations in *proS* that restore aminoacylation of G37-unmodified tRNA^Pro^ [136]. However, the locations of the *proS* mutations were not restricted to any specific domains.

This study once again highlights the importance of integrating analytical, computational, and genetic approaches to accurately define tRNA modification profiles in a given organism [17]. Encouragingly, growing recognition of the biological significance of RNA modifications— combined with improved access to analytical technologies—has led to a surge of studies across taxonomically diverse species [21,22,27,93,137–139]. These datasets now can serve as valuable anchors for bioinformatic and machine learning approaches to predict modification pathways in newly sequenced organisms.

## Data accessibility

Raw LC-MS/MS data have been deposited to the ProteomeXchange Consortium (http://proteomecentral.proteomexchange.org) via the PRIDE (http://www.ebi.ac.uk/pride/) partner repository with the dataset identifier PXD069173. Raw AlkAnilineSeq and GLORI data are available at the European Nucleotide Archive (https://www.ebi.ac.uk/ena/browser/home) under the accession number PRJEB96333.

## Declaration of AI use

Grammarly (https://www.grammarly.com) and ChatGPT version 4.o (https://chatgpt.com/) were used to edit the grammar and clarity of specific sentences.

## Supporting information

SupTables_Figures

Supdata 1

SupData2

Supdata3

Supdata4

## Acknowledgments

We thank Patricia Dos Santos and Frederic Barras for their helpful discussions, and Stefano Marzi for sharing unpublished data and providing input on the manuscript.

