## Supplementary material for "tRNA Modification Landscapes in Streptococci: Shared Losses and Clade-Specific Adaptations": SupTables_Figures

**Supplemental Material**

**Table S1. Strains and plasmids used and analytical runs using different techniques.**

| **Strains used in this study** | | | |
|---|---|---|---|
| Strain number | Genotype (description) | Antibiotic resistance[a] | Reference or source |
| **S. pneumoniae** | | | |
| IU1690 | D39 *cps*$^+$ (D39W) | None | [1,2] Duplicate tRNA preparations in [3] |
| IU1781 | D39 *cps*$^+$ *rpsL1* | Str$^R$ | [1] |
| IU1824[c] | D39 *rpsL1* Δ*cps2A'-cps2H'* = D39 *rpsL1* Δ*cps* *spd_0567*(*mnmM*)(Q152stop) | Str$^R$ | [1] One tRNA preparations in [3] |
| IU1945 | D39 Δ*cps2A'-cps2H'*= D39 Δ*cps* | None | [1] |
| E46 | IU1945 Δ*bgaA::*P$_c$-*erm* (IU1945 X fusion Δ*bgaA::*P*c-erm*) | Erm$^R$ | This study |
| E797 | IU1945 Δ*tsaE::*P$_c$-*erm* sup 3 (with suppressor mutation *asnS* [D121A]) (IU1945 X Δ*tsaE::*P*c-erm* from IU18824) | Erm$^R$ | This study |
| K797 | IU1945 Δ*tsaE*::P$_c$-*kan-rpsL* sup 4 (with suppressor mutation *asnS* [D121A]) (IU1945 X Δ*tsaE::* P*c-kan-rpsL* from IU18826) | Kan$^R$ | This study |
| IU18816 | D39 Δ*cps rpsL1* Δ*bgaA*::*spec*-P$_{Zn}$-*tsaE*$^+$ (IU1824 X fusion Δ*bgaA*::*spec*-P$_{Zn}$-*tsaE*$^+$) | Str$^R$ Spc$^R$ | This study |
| IU18824 | D39 Δ*cps rpsL1* Δ*tsaE::*P$_c$-*erm* sup 1 (with suppressor mutation *asnS* [D121A] (IU1824 X fusion Δ*tsaE::*P$_c$-*erm*) | Str$^R$ Erm$^R$ | This study |
| IU18826 | D39 Δ*cps rpsL1* Δ*tsaE*::P$_c$-*kan-rpsL* sup 2 (with suppressor mutations *asnS* [D121A] and *spd_0982*[R191H] (IU1824 X fusion Δ*tsaE::*P$_c$-*kan-rpsL*) | Kan$^R$ | This study |
| IU18886 | D39 Δ*cps rpsL1* Δ*tsaE* P$_c$-*kan-rpsL* //Δ*bgaA*::*spec*-P$_{Zn}$-*tsaE*$^+$ (IU18816 X Δ*tsaE* P$_c$-*kan-rpsL* from E797) | Kan$^R$ Spc$^R$ | This study |
| IU18963 | D39 Δ*cps rpsL1* Δ*tsaE* //Δ*bgaA::spec*-P$_{Zn}$-*tsaE*$^+$ (IU18886 X fuson Δ*tsaE)* | Str$^R$ Spc$^R$ | This study |
| IU19598 | D39 Δ*cps rpsL1* Δ*bgaA::kan*-P$_{ftsA}$-*asnS* [D121A] (IU1824 X fusion Δ*bgaA::kan*-P$_{ftsA}$-*asnS* [D121A]) | Str$^R$ Kan$^R$ | This study. Two tRNA preparations |
| IU19599 | D39 Δ*cps rpsL1* Δ*bgaA::kan*-P$_{ftsA}$-*asnS*$^+$(IU1824 X fusion Δ*bgaA::kan*-P$_{ftsA}$-*asnS*$^+$) | Str$^R$ Kan$^R$ | This study |
| IU19699 | D39 Δ*cps rpsL1* Δ*bgaA::kan*-P$_{ftsA}$-*asnS* [D121A] Δ*tsaE::*P$_c$-*erm* (IU19598 x Δ*tsaE::*P$_c$-*erm* from IU18824) | Str$^R$ Kan$^R$ Erm$^R$ | This study. Two tRNA preparations |

| | | | |
|---|---|---|---|
| IU20484 | D39 Δ*cps rpsL1* Δ*dusB1*(*spd_2016*)::P$_{c}$-*erm* (IU1824 X fusion Δ*dusB1*(*spd_2016*)::P$_{c}$-*erm*) | Str$^{R}$ Erm$^{R}$ | This study. Three tRNA preparations. |

| | | | |
|---|---|---|---|
| ***S. mutans*** | | | |
| *S. mutans* UA159 | Serotype c, ATCC 700610<br>Kelly C. Rice laboratory Isolate | | [4]<br>Duplicate tRNA preparations in [5]. One tRNA preparation in [3] |
| *S. mutans* UA159 | Serotype c, ATCC 700610<br>Z. Wen laboratory isolate | | [6] |
| JB409 | *S. mutans* UA159 Δ*tsaE*::Kan$^{R}$ | | [6] |
| MSJ1093 | *S. mutans* UA159 Δ*ytqA*::Erm$^{R}$ | Erm$^{R}$ | Triplicate tRNA preparation in [3] |
| MSJ1095 | *S. mutans* Δ*ymnmM*::Erm$^{R}$ | Erm$^{R}$ | Triplicate tRNA preparation in [3] |
| MSJ1031 | *S. mutans* UA159 Δ*queG*::Kan$^{R}$ | Kan$^{R}$ | One tRNA preparation in [5] |
| MSJ1097 | *S. mutans* Δ*ytqA mnmM*::Erm$^{R}$ | Erm$^{R}$ | Triplicate tRNA preparation in [3] |

**Table S2.** Oligonucleotides and DNA templates used in this study.

| **Primers used to construct strains** | | | |
|---|---|---|---|
| Primer | Sequence (5' to 3') | Template$^{b}$ | Amplicon Product |
| **For construction of E46 (Δ*bgaA*::P$_{c}$-*erm*)** | | | |
| P146 | TGGCCATTCATCGCTGGTCGTGCTGAAAT | D39 | 5' fragment with 60 bp of 5' *bgaA* |
| P148 | CATTATCCATTAAAAATCAAACGGATCCTATCC CACAGCAAACTTACGAATGCTATAAAC | | |
| Kan rpsL forward | TAGGATCCGTTTGATTTTTAATGGATAATG | Pc-*erm* cassette$^{c}$ | Pc-*erm* |
| Kan rpsL reverse | GGGCCCCTTTCCTTATGCTTTTG | | |
| P149 | CAAAAGCATAAGGAAAGGGGCCCGCTCTTCT AGGTTTGAGTGCAGGATTAG | D39 | 3' fragment with 60 bp of 3' *bgaA* |
| P147 | TACGCCTTCTATCATGCCTTTGATCGCCCGT | | |
| **For construction of IU18816 (Δ*bgaA*::*spec*-P$_{Zn}$-*tsaE*$^{+}$)** | | | |
| TT657 | CGCCCCAAGTTCATCACCAATGACATCAAC | IU18560$^{d}$ | 5' Δ*bgaA*::*spec*-P$_{Zn}$ |
| TT1497 | TTGCAACTCTTCTTCATTTTTTGTGTACATTAC ATCGCTTCCTCTCTATCTTCCTTGTTA | | |
| TT1498 | TAACAAGGAAGATAGAGAGGAAGCGATGTAAT GTACACAAAAAATGAAGAAGAGTTGCAA | D39 | *tsaE*$^{+}$ |
| TT1499 | AGCAACTGGTTTATGAGAAAGTAAGTTCTTTC ATACTCCATATTGAAGCTCCTCTAACAA | | |
| TT1500 | TTGTTAGAGGAGCTTCAATATGGAGTATGAAA GAACTTACTTTCTCATAAACCAGTTGCT | D39 | *bgaA'* to downstream |
| CS121 | GCTTTCTTGAGGCAATTCACTTGGTGC | | |

<table>
<tr><td colspan="4">For construction of IU18824 (ΔtsaE::P<sub>c</sub>-erm)</td></tr>
<tr><td>TT1493</td><td>CTCTTCAAATCACGTCAGCTCTATCTGCAATC TC</td><td rowspan="2">D39</td><td rowspan="2">5’ fragment with 60 bp of 5’ tsaE</td></tr>
<tr><td>TT1494</td><td>CATTATCCATTAAAAATCAAACGGATCCTATAAT AGATGGCCCAAACGCTCCCCTAAGGC</td></tr>
<tr><td>Kan rpsL forward</td><td>TAGGATCCGTTTGATTTTTAATGGATAATG</td><td rowspan="2">Pc-erm cassette[c]</td><td rowspan="2">Pc-erm</td></tr>
<tr><td>Kan rpsL reverse</td><td>GGGCCCCTTTCCTTATGCTTTTG</td></tr>
<tr><td>TT1495</td><td>AACGTCCAAAAGCATAAGGAAAGGGGCCCTT TCAGGCAAAGGGTTTGCGTGCTGAGAAAT</td><td rowspan="2">D39</td><td rowspan="2">3’ fragment with 60 bp of 3’ tsaE</td></tr>
<tr><td>TT1496</td><td>GAGCCTGACCATTCCCTGATTCAATGCGCG</td></tr>
<tr><td colspan="4">For construction of IU18826 (ΔtsaE::P<sub>c</sub>-[kan-rpsL<sup>+</sup>])</td></tr>
<tr><td>TT1493</td><td>CTCTTCAAATCACGTCAGCTCTATCTGCAATC TC</td><td rowspan="2">D39</td><td rowspan="2">5’ fragment with 60 bp of 5’ tsaE</td></tr>
<tr><td>TT1494</td><td>CATTATCCATTAAAAATCAAACGGATCCTATAAT AGATGGCCCAAACGCTCCCCTAAGGC</td></tr>
<tr><td>Kan rpsL forward</td><td>TAGGATCCGTTTGATTTTTAATGGATAATG</td><td rowspan="2">P<sub>c</sub>-[kan-rpsL<sup>+</sup>] cassette[c]</td><td rowspan="2">P<sub>c</sub>-[kan-rpsL<sup>+</sup>]</td></tr>
<tr><td>Kan rpsL reverse</td><td>GGGCCCCTTTCCTTATGCTTTTG</td></tr>
<tr><td>TT1495</td><td>AACGTCCAAAAGCATAAGGAAAGGGGCCCTT TCAGGCAAAGGGTTTGCGTGCTGAGAAAT</td><td rowspan="2">D39</td><td rowspan="2">3’ fragment with 60 bp of 3’ tsaE</td></tr>
<tr><td>TT1496</td><td>GAGCCTGACCATTCCCTGATTCAATGCGCG</td></tr>
<tr><td colspan="4">For construction of IU18963 (ΔtsaE markerless)</td></tr>
<tr><td>TT1493</td><td>CTCTTCAAATCACGTCAGCTCTATCTGCAATC TC</td><td rowspan="2">D39</td><td rowspan="2">5’ fragment with 60 bp of 5’ tsaE</td></tr>
<tr><td>TT1513</td><td>TTTCTCAGCACGCAAACCCTTTGCCTGAAATA ATAGATGGCCCAAACGCTCCCCTAAGGC</td></tr>
<tr><td>TT1514</td><td>GCCTTAGGGGAGCGTTTGGGCCATCTATTATT TCAGGCAAAGGGTTTGCGTGCTGAGAAA</td><td rowspan="2">D39</td><td rowspan="2">3’ fragment with 60 bp of 3’ tsaE</td></tr>
<tr><td>TT1496</td><td>GAGCCTGACCATTCCCTGATTCAATGCGCG</td></tr>
<tr><td colspan="4">For construction of IU19598 (ΔbgaA::kan-P<sub>ftsA</sub>-asnS [D121A])</td></tr>
<tr><td>TT657</td><td>CGCCCCAAGTTCATCACCAATGACATCAAC</td><td rowspan="2">IU9621</td><td rowspan="2">5’ ΔbgaA::kan-P<sub>ftsA</sub></td></tr>
<tr><td>TT1586</td><td>TACATCAATAATCGTTACACGTTTTGTCATTAC ATCGCTTCCTCTCTATCTTCCAAGTTT</td></tr>
<tr><td>TT1587</td><td>AACTTGGAAGATAGAGAGGAAGCGATGTAATG ACAAAACGTGTAACGATTATTGATGTAA</td><td rowspan="2">IU18824</td><td rowspan="2">asnS [D121A]</td></tr>
<tr><td>TT1588</td><td>AGCAACTGGTTTATGAGAAAGTAAGTTCTTTTA TGGTTTGATACGGTGCAACATACGTGG</td></tr>
<tr><td>TT1589</td><td>CACGTATGTTGCACCGTATCAAACCATAAAAG AACTTACTTTCTCATAAACCAGTTGCTG</td><td rowspan="2">D39</td><td rowspan="2">bgaA’ to downstream</td></tr>
<tr><td>CS121</td><td>GCTTTCTTGAGGCAATTCACTTGGTGC</td></tr>
<tr><td colspan="4">For construction of IU19599 (ΔbgaA::kan-P<sub>ftsA</sub>-asnS [D121A])</td></tr>
<tr><td>TT657</td><td>CGCCCCAAGTTCATCACCAATGACATCAAC</td><td rowspan="2">IU9621[e]</td><td rowspan="2">5’ ΔbgaA::kan-P<sub>ftsA</sub></td></tr>
<tr><td>TT1586</td><td>TACATCAATAATCGTTACACGTTTTGTCATTAC ATCGCTTCCTCTCTATCTTCCAAGTTT</td></tr>
<tr><td>TT1587</td><td>AACTTGGAAGATAGAGAGGAAGCGATGTAATG ACAAAACGTGTAACGATTATTGATGTAA</td><td>D39</td><td>asnS<sup>+</sup></td></tr>
</table>

| | | | |
|---|---|---|---|
| TT1588 | AGCAACTGGTTTATGAGAAAGTAAGTTCTTTTA TGGTTTGATACGGTGCAACATACGTGG | | |
| TT1589 | CACGTATGTTGCACCGTATCAAACCATAAAAG AACTTACTTTCTCATAAACCAGTTGCTG | D39 | *bgaA'* to downstream |
| CS121 | GCTTTCTTGAGGCAATTCACTTGGTGC | | |
| CS121 | GCTTTCTTGAGGCAATTCACTTGGTGC | | |
| **For construction of IU20484 (Δ*dusB1* (*spd_2016*)*::*$P_c$*-erm*)** | | | |
| TT1666 | GATAGCACTGAAACCGTCCGCACTGCTCAAG | D39 | 5' fragment with 60 bp of 5' *dusB* |
| TT1668 | CATTATCCATTAAAAATCAAACGGATCCTAGGT CACGCCAGCCATAGGCGCTAA | | |
| Kan rpsL forward | TAGGATCCGTTTGATTTTTAATGGATAATG | Pc-*erm* cassette[d] | Pc-*erm* |
| Kan rpsL reverse | GGGCCCCTTTCCTTATGCTTTTG | | |
| TT1669 | CAAAAGCATAAGGAAAGGGGCCCACATCTGG CGCTGCCAAACTCCGT | D39 | 3' fragment with 60 bp of 3' *dusB* |
| TT1667 | CGCTAATGGTTCAATGGCAACAGGATGGCTC C | | |
| **Amplicons used for transformation assays** | | | |
| **Δ*tsaE*::$P_c$-*erm*** | | | |
| TT1493 | CTCTTCAAATCACGTCAGCTCTATCTGCAATCTC | IU18824 | Δ*tsaE*::$P_c$-*erm* |
| TT1496 | GAGCCTGACCATTCCCTGATTCAATGCGCG | | |
| **Δ*tsaE*::$P_c$-[*kan-rpsL*$^+$]** | | | |
| TT1493 | CTCTTCAAATCACGTCAGCTCTATCTGCAATCTC | IU18826 | Δ*tsaE*::$P_c$-[*kan-rpsL*$^+$] |
| TT1496 | GAGCCTGACCATTCCCTGATTCAATGCGCG | | |
| **Δ*bgaA*::$P_c$-*erm*** | | | |
| P146 | TGGCCATTCATCGCTGGTCGTGCTGAAAT | E46 | Δ*bgaA*::$P_c$-*erm* |
| P147 | TACGCCTTCTATCATGCCTTTGATCGCCCGT | | |
| **Δ*tsaE*::markerless** | | | |
| TT1493 | CTCTTCAAATCACGTCAGCTCTATCTGCAATCTC | IU18963 | Δ*tsaE* markerless |
| TT1496 | GAGCCTGACCATTCCCTGATTCAATGCGCG | | |

[a]Antibiotic resistance markers: $Erm^R$, erythromycin; $Kan^R$, kanamycin; $Spc^R$, spectinomycin; $Str^R$, streptomycin; $Tet^R$, tetracycline.

[b]Genomic DNA of indicated *S. pneumoniae* strains was used as templates for PCR reactions, except for $P_c$-[*kan-rpsL*$^+$] and $P_c$-*erm* cassettes.

[c]$P_c$-*erm* and $P_c$-[*kan-rpsL*$^+$] cassettes are described in [7].

[d] IU18560 is D39 Δ*cps rpsL1* Δ*bgaA*::*spec*-$P_{Zn}$-*stkP*$^+$, Winkler lab stock.

[e] IU9621 is D39 Δ*cps rpsL1* Δ*khpA*//Δ*bgaA*::*kan*-$P_{ftsA}$-*khpA*$^+$ , Winkler lab stock.

**Supplemental Data 1 Gene, protein and genome data**
**1A** list of *B. subtilis* tRNA modification proteins
**1B** list of *S. mutans* UA159 tRNA modification proteins and essentiality data
**1C** list of *S. pneumoniae* D39 tRNA modification proteins and essentiality data
**1D** Compilation of tRNA modification data for the three model Gram-positive
**1E** Distribution of Q synthesis and salvage genes in 1599 Streptococci
**1F** Identifiers for sequences used in multiple alignments
**1G** Identifiers of sequences used in SSN
**1H** Gene neighborhood data
**1I** Codon usage data
**1J** Essentiality of *tsaE* in ESKAPE pathogens

**Supplemental Data 2 LC/MS analyses**
**2A** List of detected modified nucleosides confirmed by standards
**2B** LC/MS analyses 11/08/2021 and Sun 10/30/2023
**2C** LC/MS 10/30/2023
**2D** LC/MS 03/18/2025
**2E** LC/MS 4/17/2025
**2F** LC/MS 10/01/2025

**Supplemental Data 3 Supplemental LC/MS analyses**

**Supplemental Data 4. COG and KO data**
**4A** List of known tRNA modification proteins with associated COG and KO family numbers
**4B** Presence/absence of tRNA modification COGs in 4 model Streptococci
**4C** Presence/absence of tRNA modification KO in 4 model Streptococci

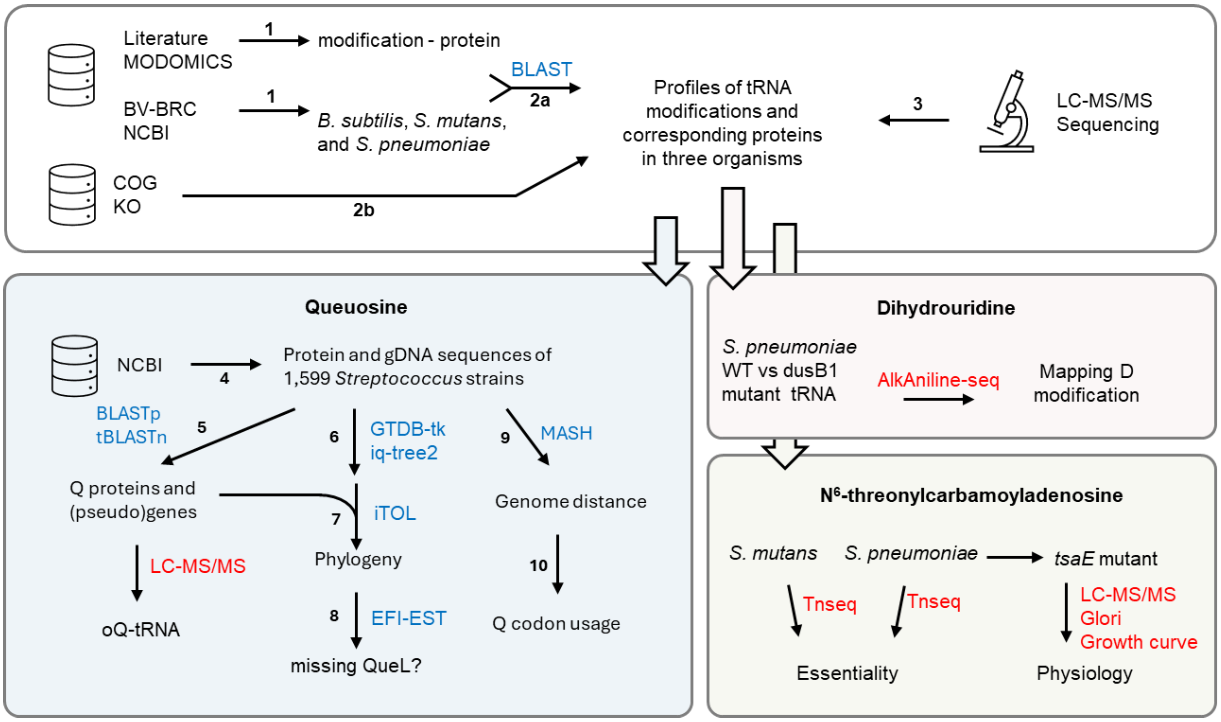


**Figure S1. Study workflow.** Steps of experiments and analyses (arrows) are described in the Method section. The experiments conducted and tools used are indicated in red and blue, respectively.

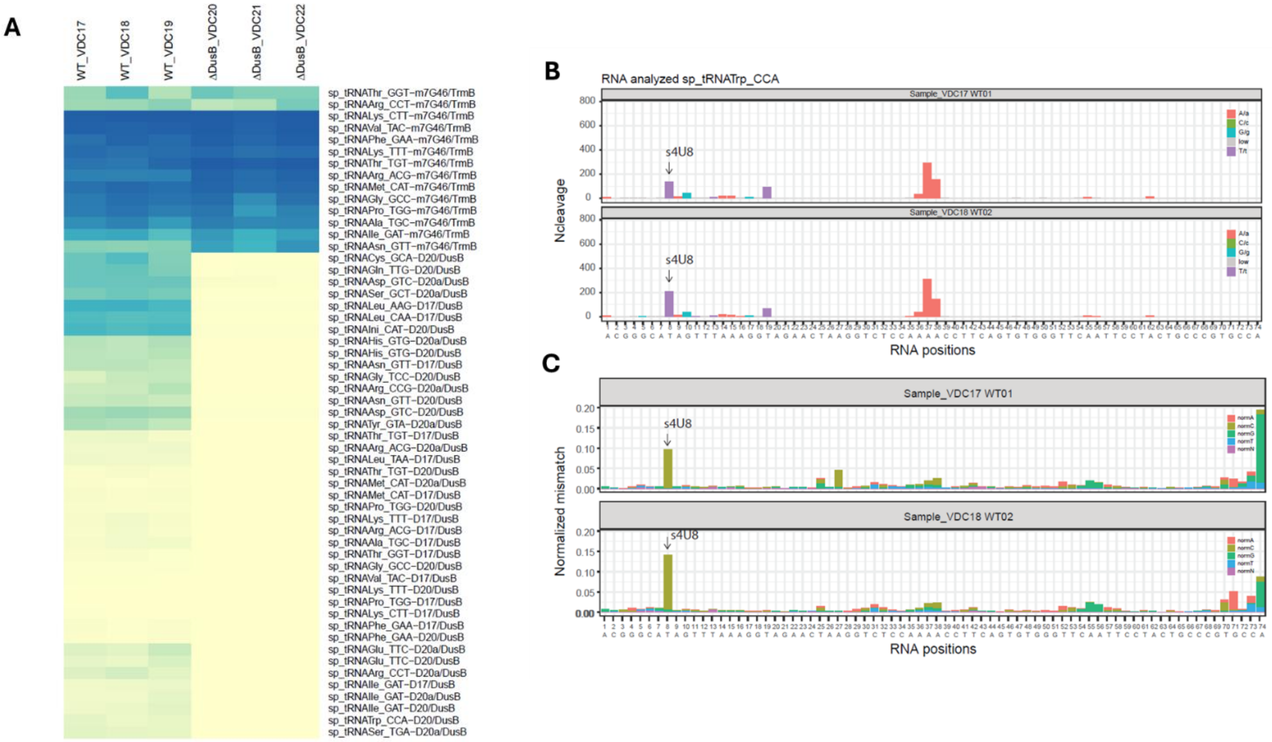


**Figure S2. AlkAnilineSeq analysis of D residues in *S. pneumoniae* tRNAs**. **(A)** Heatmap shows absolute Ncleavage values for all modified D and m$^7$G sites mapped in *S. pneumoniae* WT and D*dusB1* strains. Color code is shown at the bottom right, identity of the D site in tRNA is indicated. Data are shown for all replicates (n=3) for WT and Δ*dusB1* strains; **(B)** Two upper panels show Ncleavage signals obtained for representative tRNA (tRNATrp_CCA). Two WT tRNA samples are shown. Identity of nucleotides is shown on the right, tRNA positions are numbered in a sequential order. Position of s$^4$U8 is shown by an arrow; **(C)** The corresponding traces of normalized mismatch for the same tRNA (tRNATrp_CCA), data are extracted from the AlkAnilineSeq dataset. Color code corresponds to the nucleotide identity found at a given position in the sequence (normA, C, G, T). tRNA positions are numbered in a sequential order. Position of s$^4$U8 showing characteristic U->C transitions is shown by an arrow.

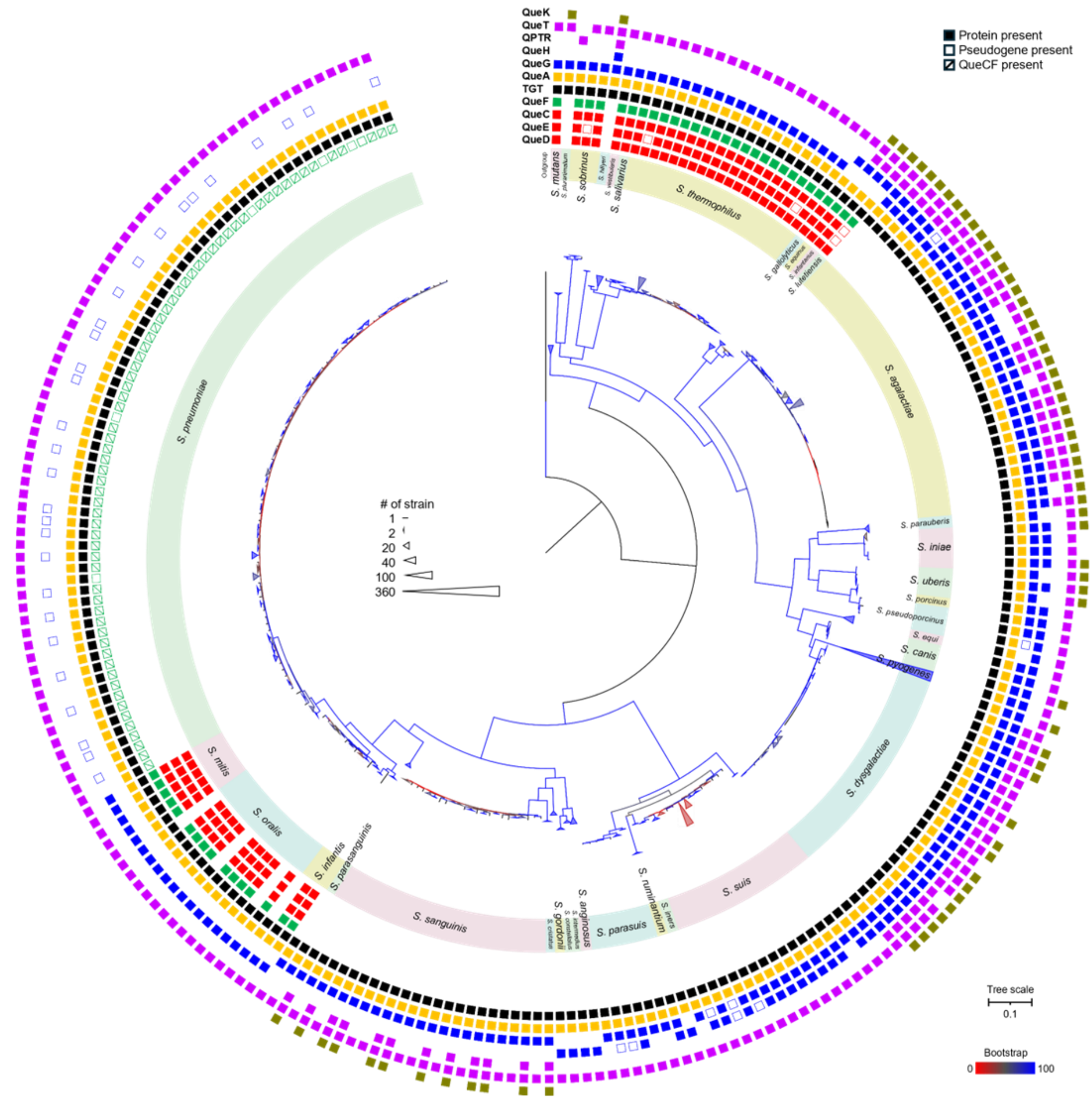


**Figure S3. Distribution of Q proteins in Streptococcus.** The maximum likelihood tree of 1,599 *Streptococcus* strains was generated based on the aligned GTDB marker proteins. Strains from *Pseudolactococcus* and *Lactococcus* were used as the outgroup. The color of the branches indicates the level of bootstrap support. Species are labeled in the inner circle. The filled and open squares in the outer circle represent the presence of corresponding Q proteins and pseudogenes, respectively. For better visualization, branches representing strains of the same species that carry identical Q gene patterns are collapsed as presented in triangles. The number of strains in each collapsed node corresponds to the size of the triangle, as shown in the legend. Species represented by fewer than two strains or rare Q gene patterns are hidden, as described in the Method section and Data S1E.

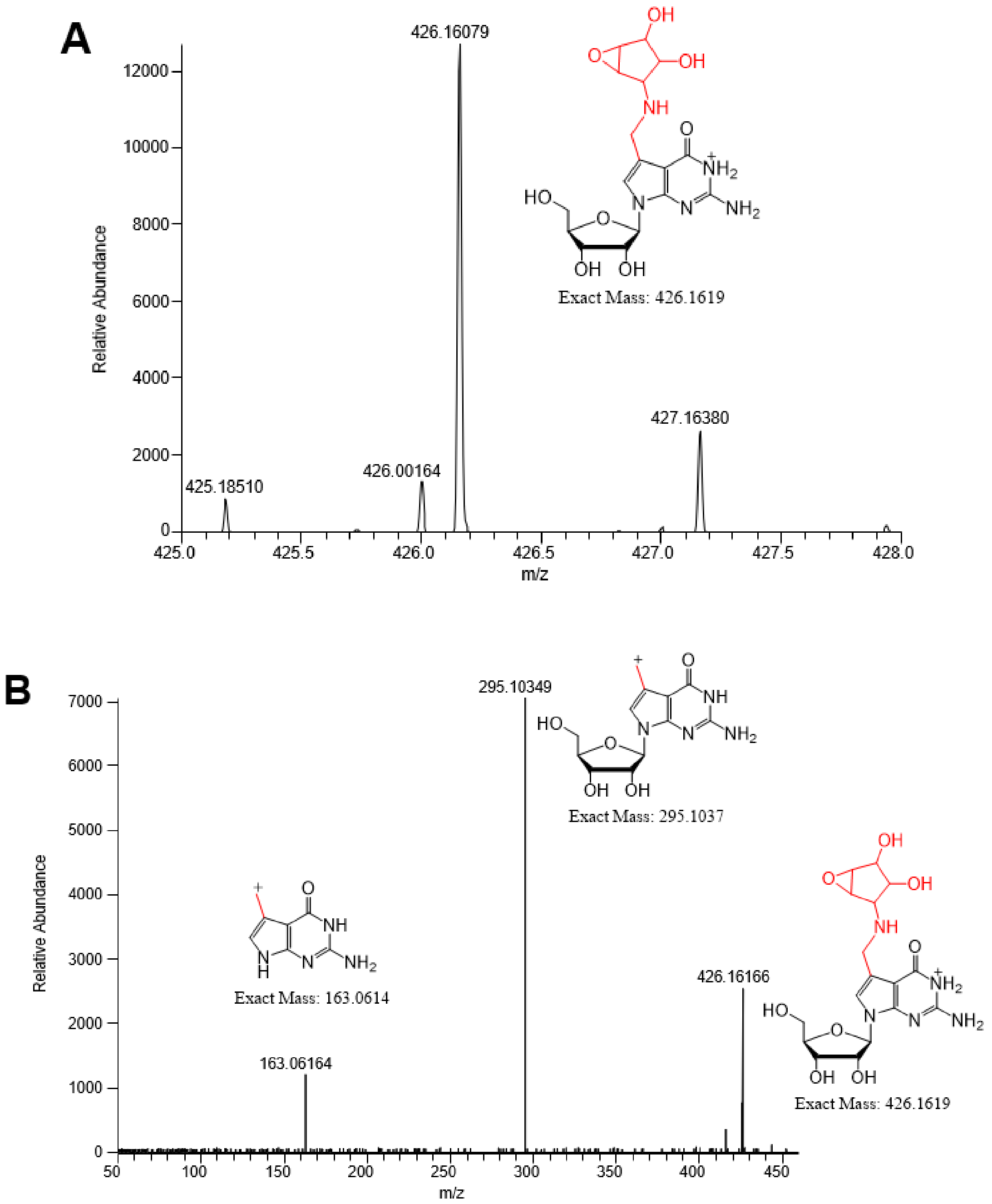


**Figure S4. High-resolution LC-MS analysis of oQ in *S. pneumoniae* tRNAs. (A)** Targeted selected ion monitoring spectrum of the ion at *m/z* 426.1619, corresponding to oQ, detected in hydrolyzed *S. pneumoniae* tRNAs; **(B)** MS/MS spectrum obtained from collision-induced dissociation of the ion at *m/z* 426.1619. The fragment ions at *m/z* 295.1035 and 163.0616 correspond to the 7-methyl-7-deazaguanosine and 7-methyl-7-deazaguanine moieties of oQ, respectively, confirming the identity of oQ.

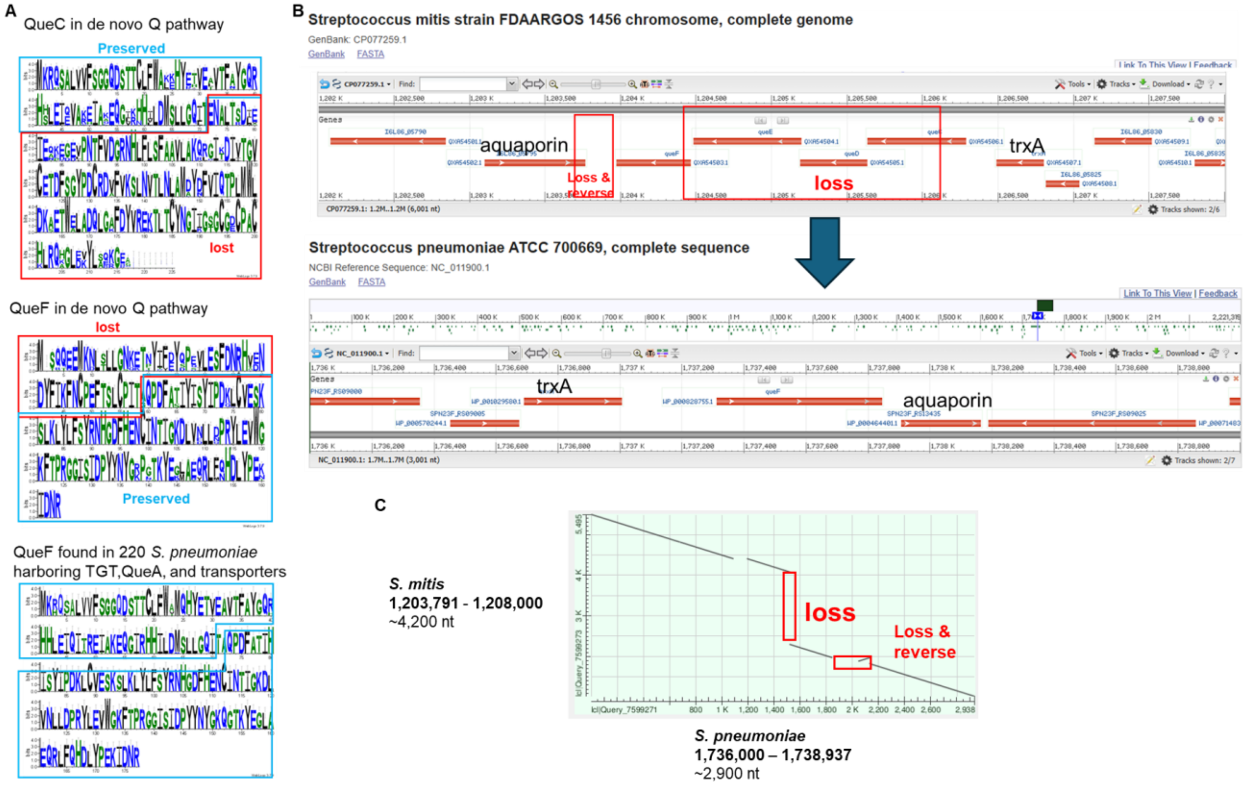


**Figure S5. QueCF fusion proteins could be the result of gene loss event**. **(A)** The sequence logos of QueC and QueF proteins from strains with the de novo Q pathway, and that of QueF proteins from 220 *S. pneumoniae* strains that encode TGT, QueA and QptR/QueT proteins. The blue and red boxes indicate the regions preserved and lost, respectively. The two preserved regions concatenate and generate the QueCF fusion proteins; **(B)** The possible gene loss from queC's C terminal, queDE, and queF's N terminal; **(C)** The alignment of genomic regions in (B). Sequences used in the alignment are listed in Supplementary data 1F.

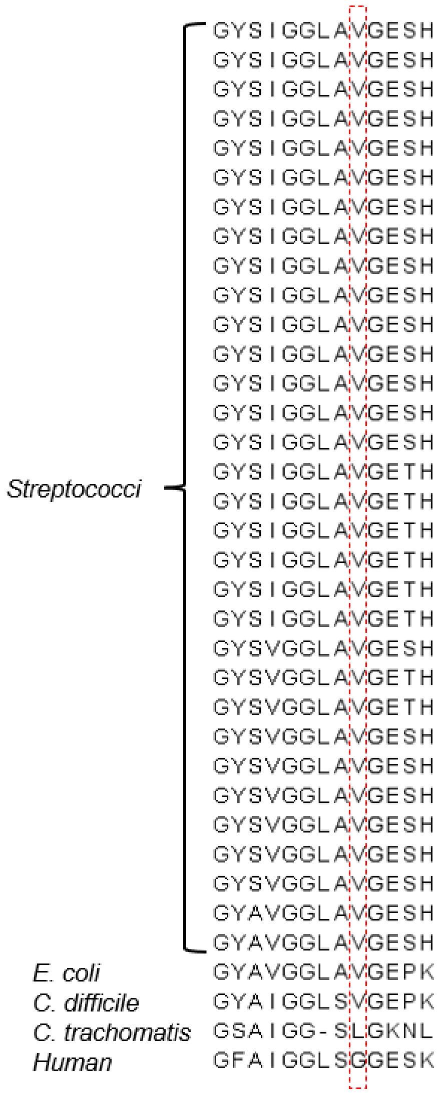


**Figure S6. The substrate binding pockets of TGTs from *Streptococci* resemble that of TGT from *E. coli***. The region of substrate binding aligned unique TGT sequences from 1,599 *Streptococcus* strains and TGTs from *E. coli, C. difficile, C. trachomatis,* and human of which the substrates are $preQ_1$, $preQ_1$, q, and q, respectively. For better visualization, only 20 TGT sequences from *Streptococcus* are shown. The boxed residue was predicted to play a pivotal role in substrate binding. Sequences used in the alignment are listed in Supplementary data 1F.

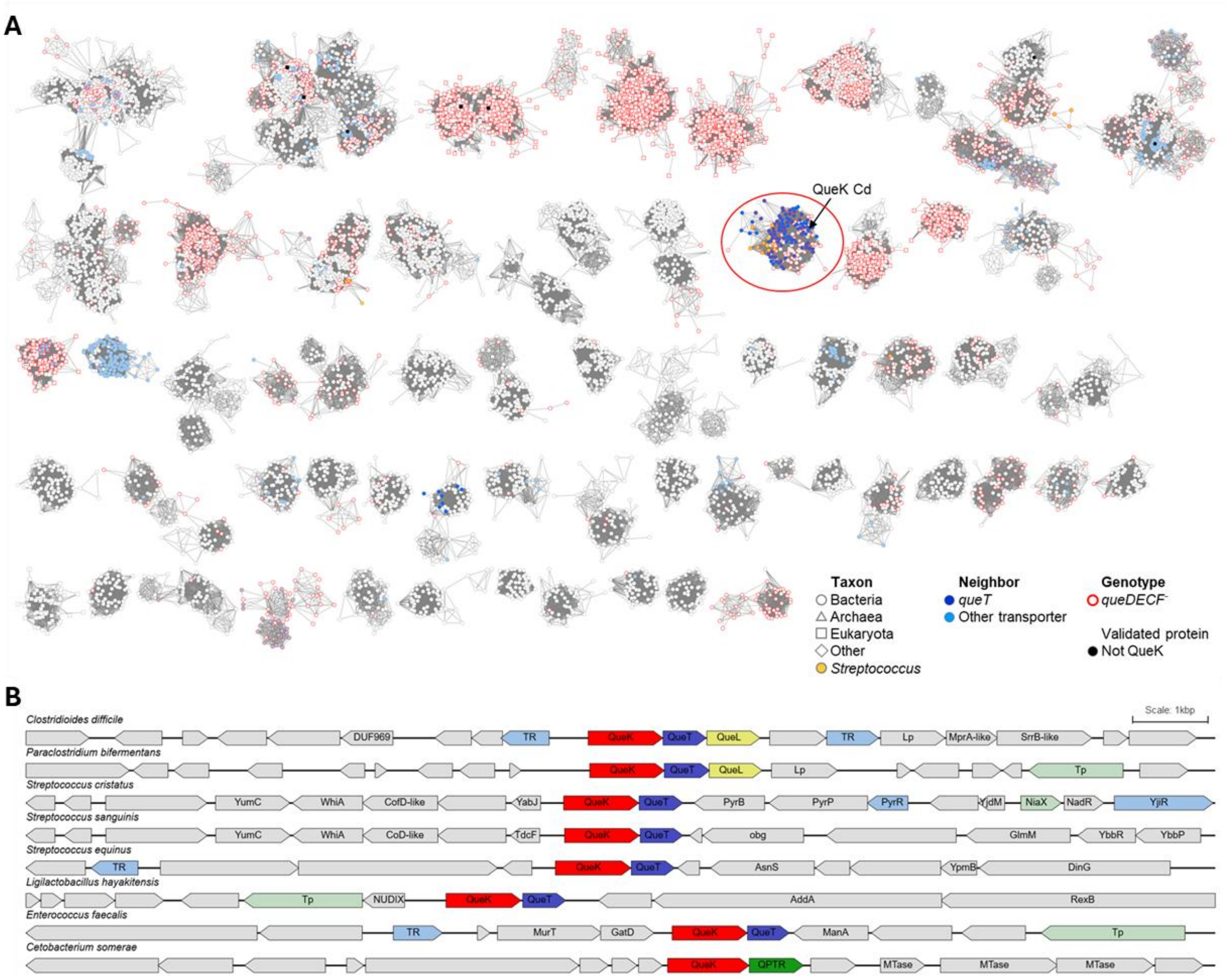


**Figure S7.** Identification of QueK protein subgroup in the IPR023186 SSN. **(A)** The SSN of Inosine/uridine-preferring nucleoside hydrolase (IPR023186) family. Each node in the network represents one or multiple sequences that share 90% identity and more. An edge (a line) is drawn between two nodes with a BLAST E-value cutoff better than $10^{-100}$ (alignment score threshold of 100). The shapes of nodes are based on the kingdom of species of the representative sequence of each node. The nodes of *Streptococcus* are filled in yellow. The nodes that have been annotated experimentally as not being QueK are filled in black. The nodes are filled in blue and skyblue when their encoding gene is next to *queT* or other transporter gene, respectively. The borders of nodes are highlighted in red when the representative sequences of each node from a genome lacks *queDECF* genes. For better visualization, clusters with less than 50 nodes are hidden; **(B)** The *queK* genes in *Streptococcus* are next to *queT* genes. The example organisms were selected from the circled cluster in the SSN. Abbraviateions: TR, putative transcription regulator; Lp, lipoprotein; Tp, transporter. The genomic information is listed in Supplementary data 1H.

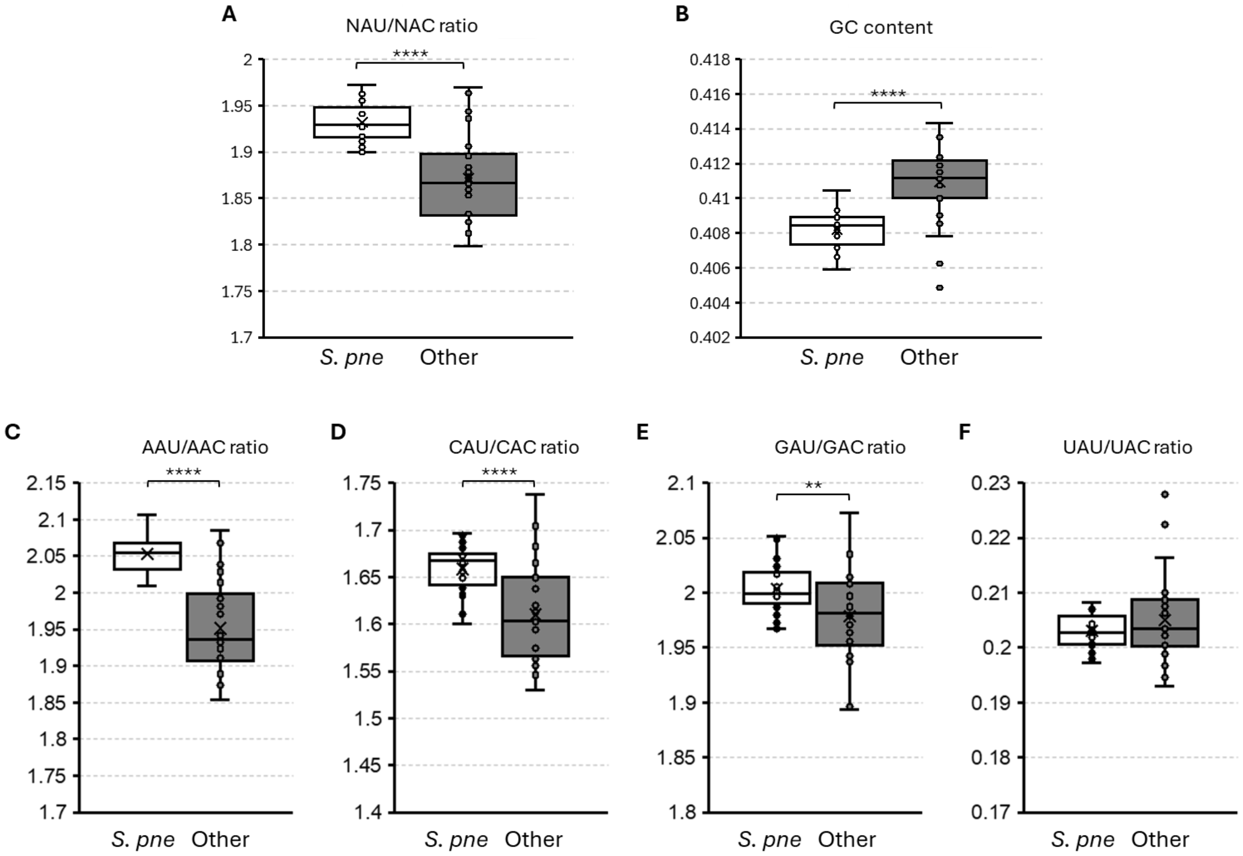


**Figure S8. Comparison of codon usage in 26 paired strains. (A)** The comparison of NAU to NAC ratio; (**B)** The comparison of GC content; **(C)** to (**F)** The comparison of AAU to AAC ratio, CAU to CAC ratio, GAU to GAC ratio, and UAU to UAC ratio, respectively. *S. pne* stands for *S. pneumoniae*. Other indicates other *Streptococcus* species. P-value of paired two-tail t.test was calculated. **: P- value < 0.01. ****: P- value < 0.0001.

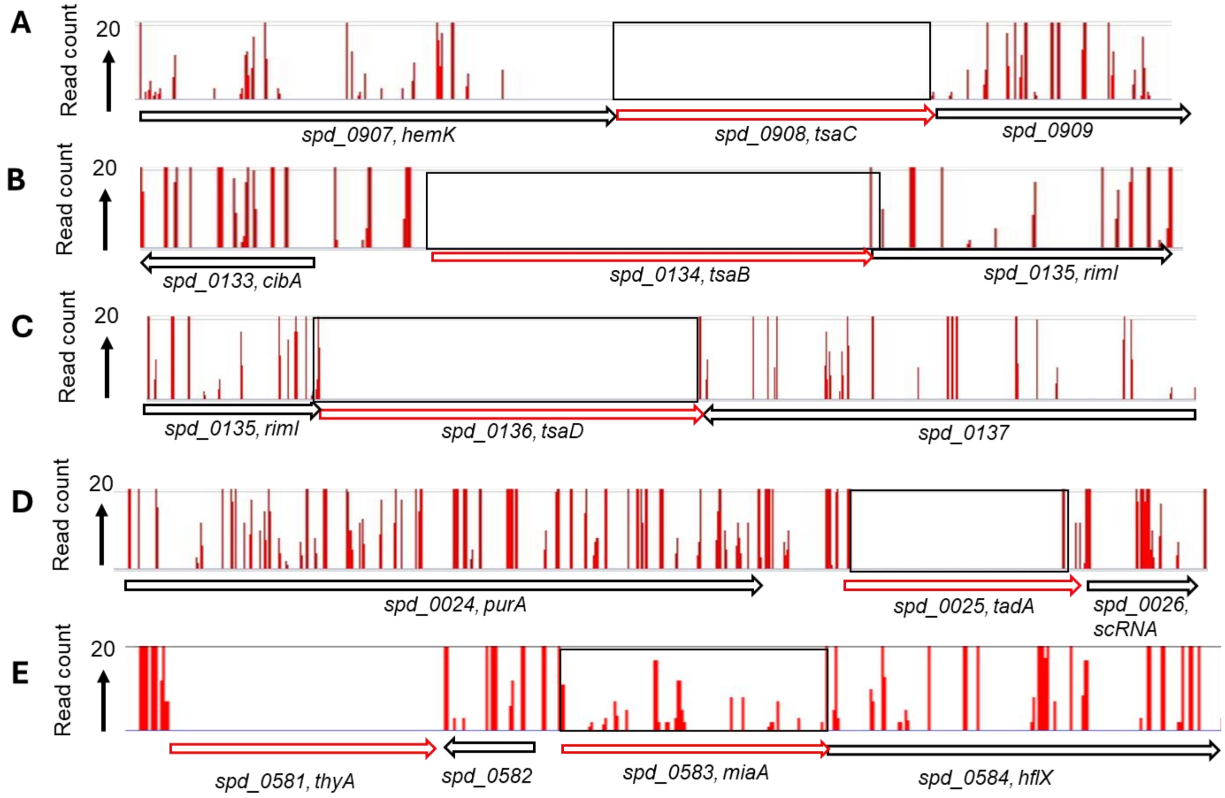


**Figure S9. TsaC, TsaB, TsaD, TadA, but not MiaA are essential in *S. pneumoniae* D39**. Mini-Mariner Malgellan6 Tn-Seq transposon insertion profile for the genome region covering **(A)** *tsaC*, **(B)** *tsaB*, **(C)** *tsaD*, **(D)** *tadA* and **(E)** *miaA* in the genomes of the unencapsulated WT parent (D39 Δcps rpsL1, IU1824).

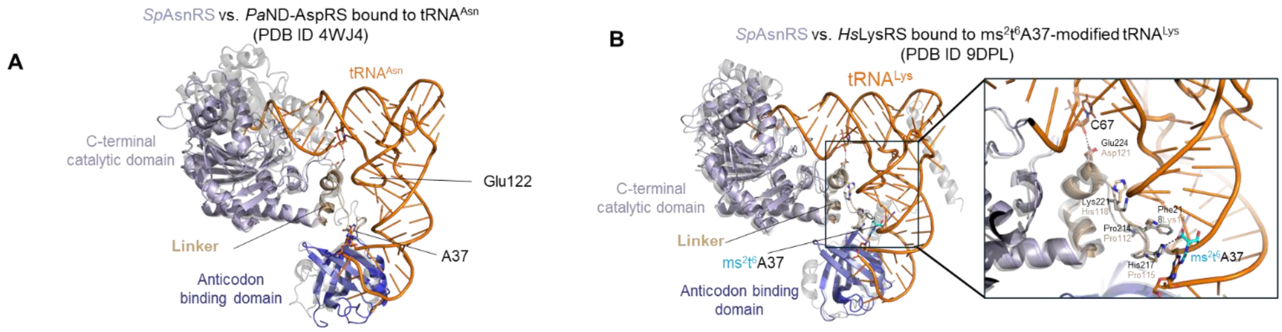


**C**

```
SptRNA-Asn-GTT-1-1      UCCGCCAUAGCUCAGUUGGUAGUAGCGCAUGACUGUUAAUCAUGAUGUCGUAGGUUCGAG 60
SmtRNA-Asn-GTT-1-1      UCCGGCAUAGCUCAGUUGGUAGUAGCGCAUGACUGUUAAUCAUGAUGUCGUAGGUUCGAG 60
                        **** *******************************************************

SptRNA-Asn-GTT-1-1      UCCUACUGGCGGAG          74
SmtRNA-Asn-GTT-1-1      UCCUACUGCCGGAG          74
                        ******** *****
```

**Figure S10**. **Structural similarity of *Sp*AsnRS with two class IIb aminoacyl-tRNA synthetases of known crystal structures bound to cognate tRNAs. (A)** AlphaFold model of *Streptococcus pneumoniae* asparaginyl-tRNA synthetase (*Sp*AsnRS, purple) superposed onto the crystal structure of *P. aeruginosa* non-discriminating aspartyl-tRNA synthetase (*Pa*ND-AspRS, grey) bound to its substrate $tRNA^{Asn}$. residues are from *Pa*ND-AspRS; **(B)** Superposition of *Sp*AsnR (purple) with *Homo sapiens* lysyl-tRNA synthetase (*Hs*LysRS, grey) bound to $ms^2t^6A37$-modified $tRNA^{Lys}$ . Linker residues interacting with tRNA in LysRS and the corresponding residues in *Sp*AsnRS are shown in stick representation and labeled. **(C)** Alignment of *S. pneumoniae* and *S. mutans* $tRNA^{Asn}$ sequences showing the one base pair difference.

| Predicted mutations | | | | | |
|---|---|---|---|---|---|
| evidence | position | mutation | annotation | gene | description |
| RA | 9,729 | A→G | Y33C (TAT→TGT) | *SMU_12* → | conserved hypothetical protein |
| RA | 11,066 | G→A | R75Q (CGA→CAA) | *SMU_13* → | putative cell-cycle protein |
| RA | 294,438 | C→T | R258C (CGT→TGT) | *pgi* → | glucose-6-phosphate isomerase |
| RA | 294,544 | A→G | E293G (GAA→GGA) | *pgi* → | glucose-6-phosphate isomerase |
| RA | 385,786 | C→T | S370F (TCT→TTT) | *brpA* → | putative transcriptional regulator |
| RA | 418,392 | T→C | F59S (TTT→TCT) | *SMU_448* → | hypothetical protein |
| RA | 552,450 | Δ1 bp | coding (395/477 nt) | *furR* → | putative ferric uptake regulator protein FurR |
| RA | 808,802 | C→G | A422A (GCC→GCG) | *pyrAB* → | carbamoylphosphate synthetase, large subunit |
| RA | 826,253 | C→G | D72H (GAC→CAC) | *SMU_875c* ← | putative transposase, IS150-like |
| RA | 842,678 | G→T | V224V (GTG→GTT) | *galE* → | UDP-galactose 4-epimerase, GalE |
| RA | 988,399 | T→G | intergenic (-8/+8) | *SMU_1039c* ← / ← *SMU_1040c* | putative lipopolysaccharide glycosyltransferase/putative oxidoreductase, short-chain dehydrogenase/reductase |
| RA | 1,164,137 | C→G | V48V (GTG→GTC) | *pyrF* ← | putative orotidine-5'-decarboxylase PyrF |
| RA | 1,223,786 | T→C | L219L (TTA→CTA) | *SMU_1297* → | conserved hypothetical protein |
| RA | 1,380,458 | C→A | G54G (GGC→GGA) | *SMU_1450* → | putative amino acid permease |
| RA | 1,423,977 | T→G | K520N (AAA→AAC) | *rexA* ← | putative exonuclease RexA |
| RA | 1,568,890 | C→T | intergenic (-93/+26) | *SMU_1647c* ← / ← *SMU_1648c* | putative transcriptional regulator/hypothetical protein |
| RA | 1,576,987 | G→C | R32G (CGC→GGC) | *nrgA* ← | putative ammonium transporter, NrgA protein |
| JC | 1,892,336 | $(TAA)_{4\rightarrow 2}$ | coding (806-811/840 nt) | *SMU_2167* ← | 50S Ribosomal Protein L2 |
| RA | 1,932,168 | $(T)_{6\rightarrow 7}$ | intergenic (-22/-133) | *SMU_2057c* ← / → *SMU_2058* | putative cadmium-transporting ATPase; P-type ATPase/putative transcriptional regulator |

**Figure S11.** Comparison of *S. mutans* WT and *tsaE* strains. Sequencing of an isogenic WT/Δ*tsaE* *S. mutans* compared to Reference Genome sequence (AE014133). The mutation boxed in red is the only one present only in the Δ*tsaE* strain.
