## Supplementary material for "tRNA Modification Landscapes in Streptococci: Shared Losses and Clade-Specific Adaptations": Supdata3

**Sup Data 3 Supplemental LC-MS analyses**

*Analysis 1. Differentiation of m^3^C, m^4^C, and m^5^C in Streptococcus pneumoniae (IU1824) tRNA hydrolysates*

m^3^C, m^4^C, and m^5^C are isobaric positional isomers that exhibit highly similar chromatographic behavior and are typically monitored using an identical MRM transition (*m/z* 258 → 126) in LC-MS/MS analyses, which complicates their accurate identification. The analytical column and LC-MS/MS method we used for the initial analysis, as detailed in the **Methods** section, were able to achieve baseline separation of m^3^C and m^5^C and detect both ribonucleosides across all tRNA samples (**Sup Data 2**). However, m^4^C was neither considered nor acquired for data during this analysis. The purpose of this supplemental analysis is to include m^4^C in the analysis and to re-confirm the peak identities of these three isomers in the tRNA samples.

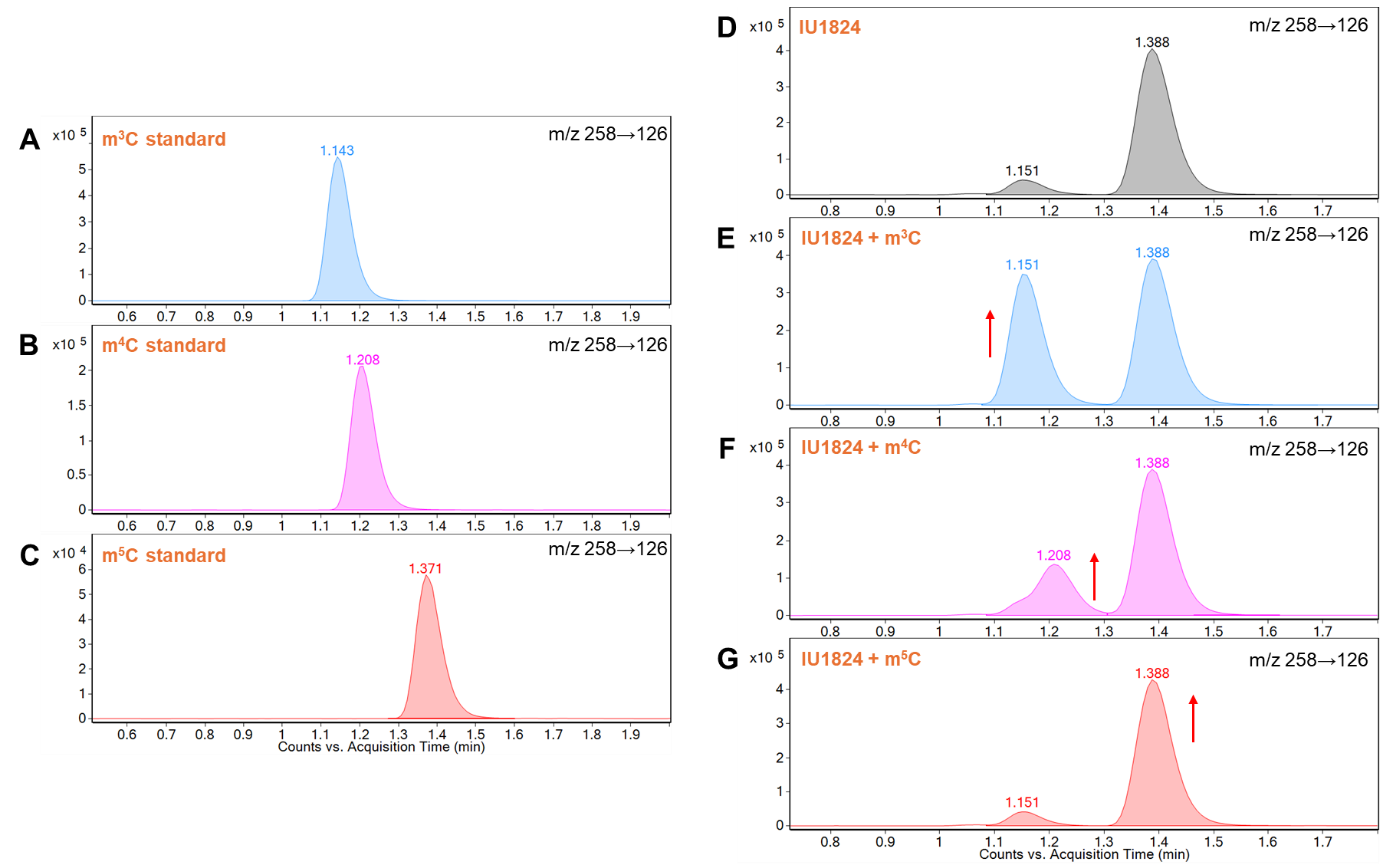

**Figure 1.** Extracted ion chromatograms obtained from LC-MS/MS analysis of (A-C) synthetic standards of m^3^C, m^4^C, and m^5^C, (D) tRNA hydrolysate of *S. pneumoniae* (IU1824), and (E-G) tRNA hydrolysate spiked with synthetic standards of m^3^C, m^4^C, and m^5^C using a Waters ACQUITY UPLC BEH C18 column (50 × 2.1 mm i.d., 1.7 µm), mobile phases, and LC gradient detailed in the **Methods** section.

As shown in **Figures 1A-C**, m^5^C was well separated from m^3^C and m^4^C, which were only partially resolved under the LC condition detailed in the **Methods** section. Analysis of *S. pneumoniae* tRNA hydrolysate detected two peaks in the MRM transition window of *m/z* 258 → 126, a typical MRM transition common to all three isomers (**Figure 1D**). By comparing retention times with synthetic standards, we first assigned the peaks at 1.15 and 1.39 min as m^3^C and m^5^C, respectively. Because the observed incomplete chromatographic separation could allow a low abundance of m^4^C to co-elute with m^3^C, we further validated peak identities through spiking individual standards to the hydrolysate (**Figure 1E-G**) and comparing peak signal intensities in unspiked and spiked samples. These results confirmed the presence of m^5^C in the samples, while the detection of m^3^C and/or m^4^C remains inconclusive and requires further investigation.

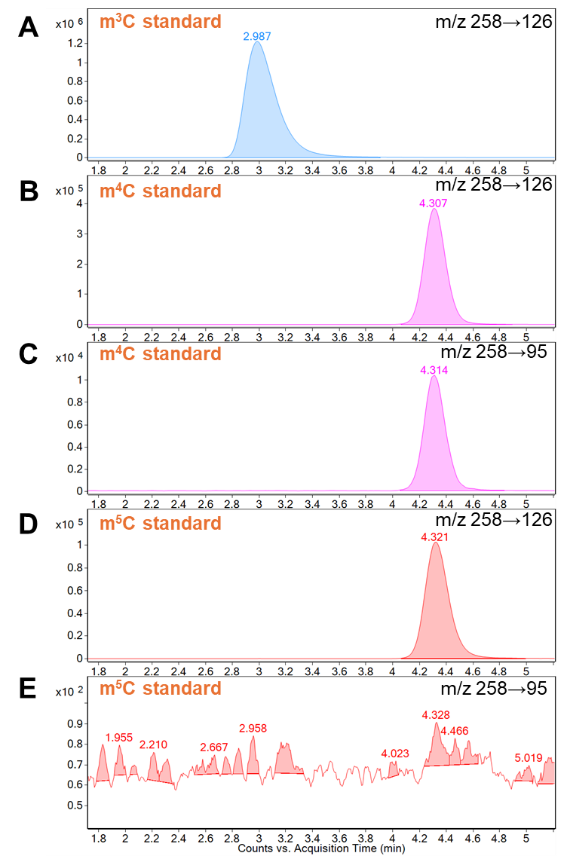

**Figure 2.** Extracted ion chromatograms obtained from LC-MS/MS analysis of synthetic standards of (A) m^3^C, (B,C) m^4^C, and (D,E) m^5^C using a Waters ACQUITY UPLC HSS T3 column (100 mm × 1 mm i.d., 1.8 µm) eluted with 5 mM ammonium acetate in water (pH 5.25) at a flow rate of 0.1 mL/min for 7 min.

To achieve baseline separation of m^3^C and m^4^C, we employed a Waters ACQUITY UPLC HSS T3 column (100 mm × 1 mm i.d., 1.8 µm) and adjusted the pH of the aqueous mobile phase to pH 5.25. The pH adjustment was intended to modulate the ionization states of the three isomers according to their reported pKa values (8.7 for m^3^C, 4.6 for m^4^C, and 4.9 for m^5^C) [1]. At pH 5.25, m^3^C exists predominately in its protonated form, whereas m^4^C and m^5^C remain largely neutral, a difference that may facilitate their chromatographic separation. The column was eluted with 5 mM ammonium acetate in water (Buffer A, pH 5.25) at a flow rate of 0.1 mL/min for 7 min, followed by a 5-min flush with pure acetonitrile (Buffer B) and an 8-min re-equilibration with 100% Buffer A prior to the next injection. As depicted in **Figures 2A and 2B,** these changes successfully achieved baseline separation of m^3^C and m^4^C, while causing coelution of m^4^C and m^5^C (**Figures 2B and 2D**). To address this limitation, we compared the published higher-energy collisional dissociation (HCD) fragmentation patterns of m^4^C and m^5^C and identified the product ion at *m/z* 95 as unique to m^4^C [2]. We then generated collision-induced dissociation fragmentation spectra of m^4^C and m^5^C on our mass spectrometer (**Figure 3**), which closely aligned with the published spectra. Therefore, we incorporated an additional MRM transition (*m/z* 258 → 95) to distinguish m^4^C and m^5^C (**Figures 2C and 2E**), where no peak was observed for m^5^C.

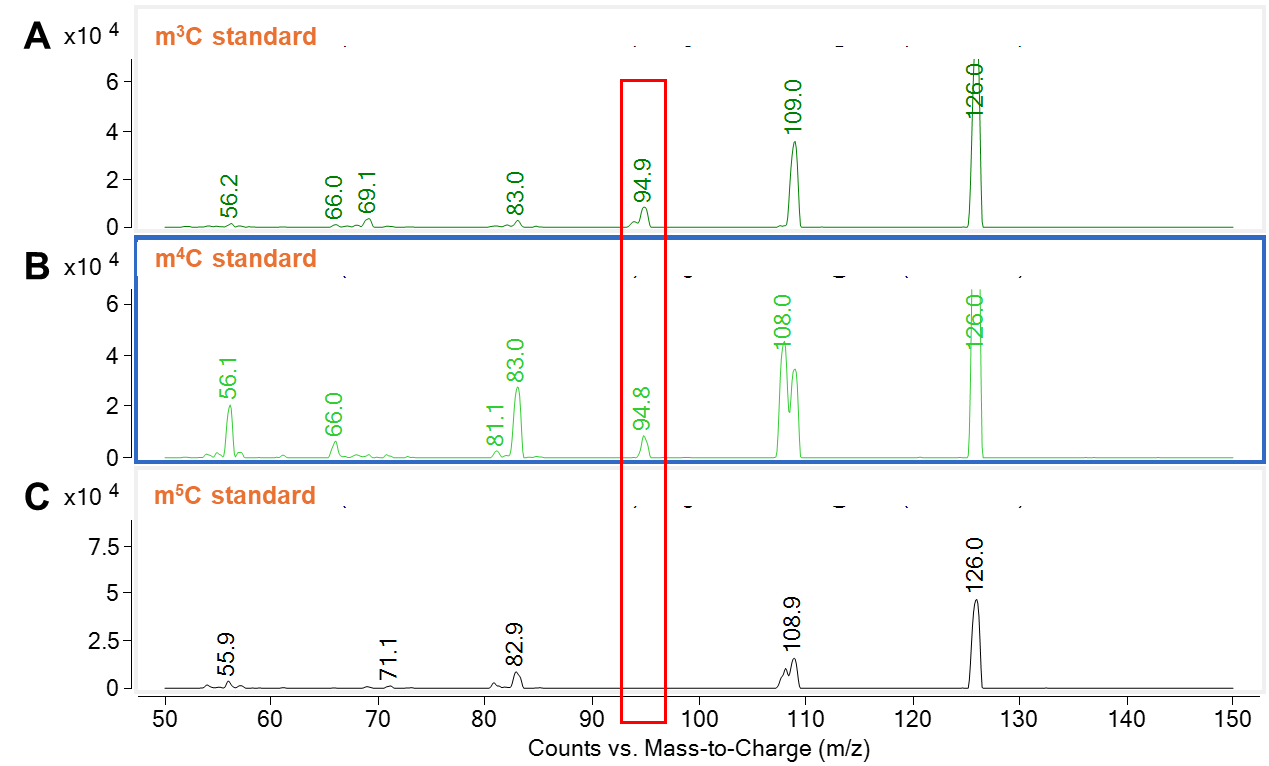

**Figure 3.** MS/MS spectra obtained from collision-induced dissociation of the protonated ions of (A) m^3^C, (B) m^4^C, and (D) m^5^C at *m/z* 258 using a collision energy of 40V.

With the ability to chromatographically resolve m^3^C and m^4^C and to distinguish m^4^C from m^5^C using a second MRM transition, we applied this method to re-analyze the *S. pneumoniae* tRNA hydrolysate and the hydrolysate spiked with individual standards. **Figures 4A-D** and **Figures 4E-H** depict the chromatograms obtained by monitoring eluates with two MRM transitions, *m/z* 258 → 126 and *m/z* 258 → 95, respectively. Comparison of peak signal intensities between unspiked and spiked samples indicated the presence of both m^3^C and m^5^C in the *S. pneumoniae* tRNA sample, whereas m^4^C was either absent or present at levels below the limit of detection.

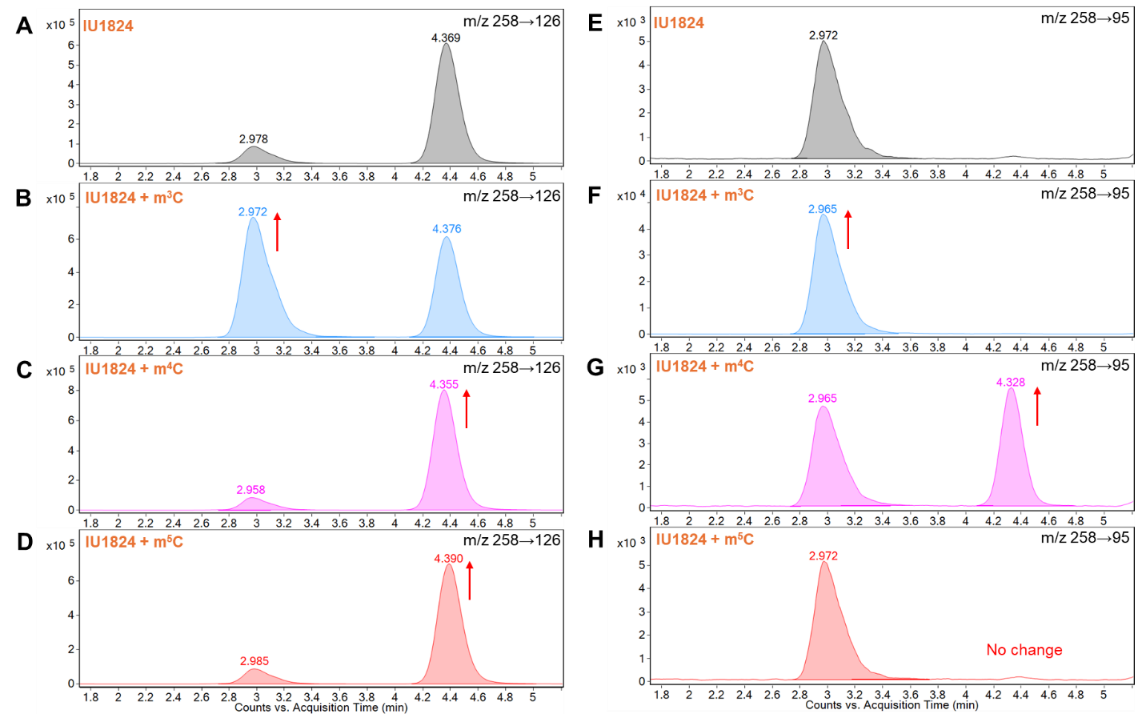

**Figure 4.** Extracted ion chromatograms obtained from LC-MS/MS analysis of (A,E) tRNA hydrolysate of *S. pneumoniae* (IU1824) and tRNA hydrolysate spiked with synthetic standards of (B,F) m^3^C, (C,G) m^4^C, and (D,H) m^5^C using a Waters ACQUITY UPLC HSS T3 column (100 mm × 1 mm i.d., 1.8 µm) eluted with 5 mM ammonium acetate in water (pH 5.25) at a flow rate of 0.1 mL/min for 7 min.

*Analysis 2. Detection of m^4^Cm in Streptococcus pneumoniae (IU1824) tRNA hydrolysates*

The absence of m^4^C in the *S. pneumoniae* tRNA sample was inconsistent with the *in-silico* data, which identified homologs of RsmH that generally catalyzes the insertion of m^4^C into rRNA. This prompted us to test for the presence of m^4^Cm, which was previously detected in the 16S rRNA of *E. coli* [3]. Because a synthetic standard of m^4^Cm was not available in our inventory, we used m^5^Cm as a reference analyte to help locate candidate peaks for m^4^Cm. With reference to the chromatographic profiles of m^3^Cm, m^4^Cm, and m^5^Cm on a C18 column by Cheng *et al.* [4], we expected m^4^Cm to elute first, followed by m^3^Cm and m^5^Cm under the LC conditions detailed in **Methods** section. **Figures 5A-C** show the chromatograms obtained from analysis of the m^5^Cm standard, the *S. pneumoniae* tRNA hydrolysate, and the hydrolysate spiked with m^5^Cm, respectively. The analysis revealed a low level of m^5^Cm (peak at 4.3 min) and a new peak at 3.4 min (**Figure 5B**). As a previous study reported that the HCD fragmentation pattern of m^4^Cm mirrors that of m^4^C [2], we incorporated multiple MRM transitions based on product ions of m^4^C to assess potential m^4^Cm peaks in the tRNA hydrolysate (**Figures 5D-H**). Given that peaks were consistently detected at the same retention times across all MRM transitions and considering the reported absence of m^3^Cm in *E. coli.* total RNA and 16s rRNA [3,4], we tentatively assigned the peak at 3.4 min as m^4^Cm. These findings suggest that m^4^C may undergo further methylation to form m^4^Cm in *S. pneumoniae* rRNA.

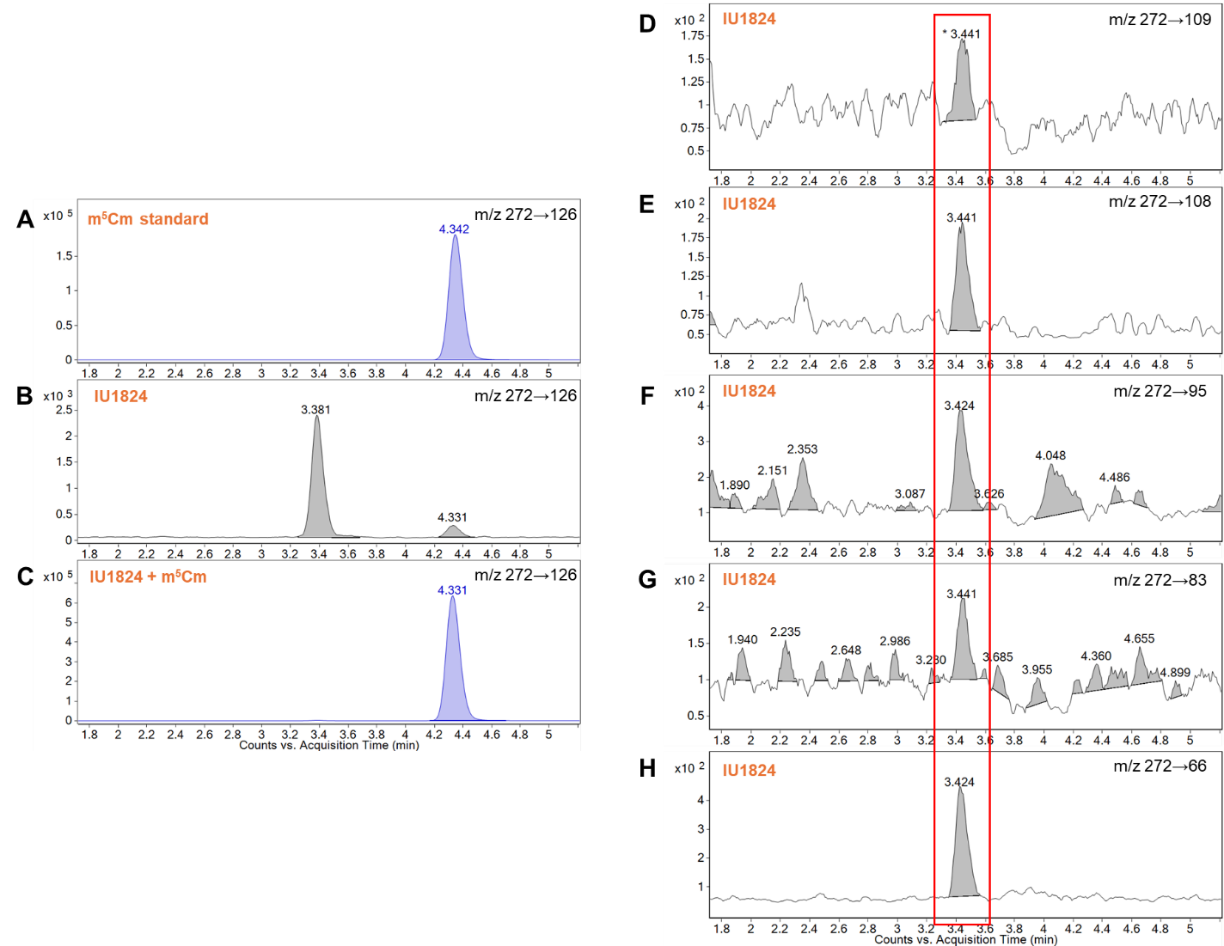

**Figure 5.** Extracted ion chromatograms obtained from LC-MS/MS analysis of (A) a synthetic standard of m^5^Cm, (B) tRNA hydrolysate of *S. pneumoniae* (IU1824), and (C) tRNA hydrolysate spiked with the synthetic standard of m^5^Cm using a Waters ACQUITY UPLC BEH C18 column (50 × 2.1 mm i.d., 1.7 µm), mobile phases, LC gradient detailed in the **Methods** section. (D-H) Additional MRM transitions were used to tentatively assign peaks at ~3.4 min as m^4^Cm.

*Analysis 3. Detection of m^4^Cm in Streptococcus mutans UA159 tRNA and rRNA* *hydrolysates*

To confirm the presence of m^4^Cm in *Streptococci* rRNA, we isolated tRNAs and rRNAs from *S. mutans* UA159, digested and analyzed with LC-MS/MS, and compared the relative abundance of m^4^Cm in the two fractions. **Figures 6A and 6B** depict the chromatograms acquired from analysis of tRNA and rRNA hydrolysate of *S. mutans* UA159, respectively, each showing a peak at ~3.5 min, consistent with that observed in **Figure 5B**. We then characterized the peak using collision-induced dissociation of the ion at *m/z* 272 (the expected precursor ion of m^4^Cm) using a series of collision energies (**Figures 6C-E**). The fragmentation pattern produced at 60 V (**Figure 6E**) is in excellent agreement with the HCD fragmentation pattern of m^4^Cm previously reported [2], confirming the peak identity as m^4^Cm. Quantitative analysis further revealed that m^4^Cm is ~750-fold more abundant in rRNA than in tRNA of *S. mutans* UA159 (**Supplemental Data 2F**), indicating that m^4^Cm is predominantly present in *Streptococci* rRNA.

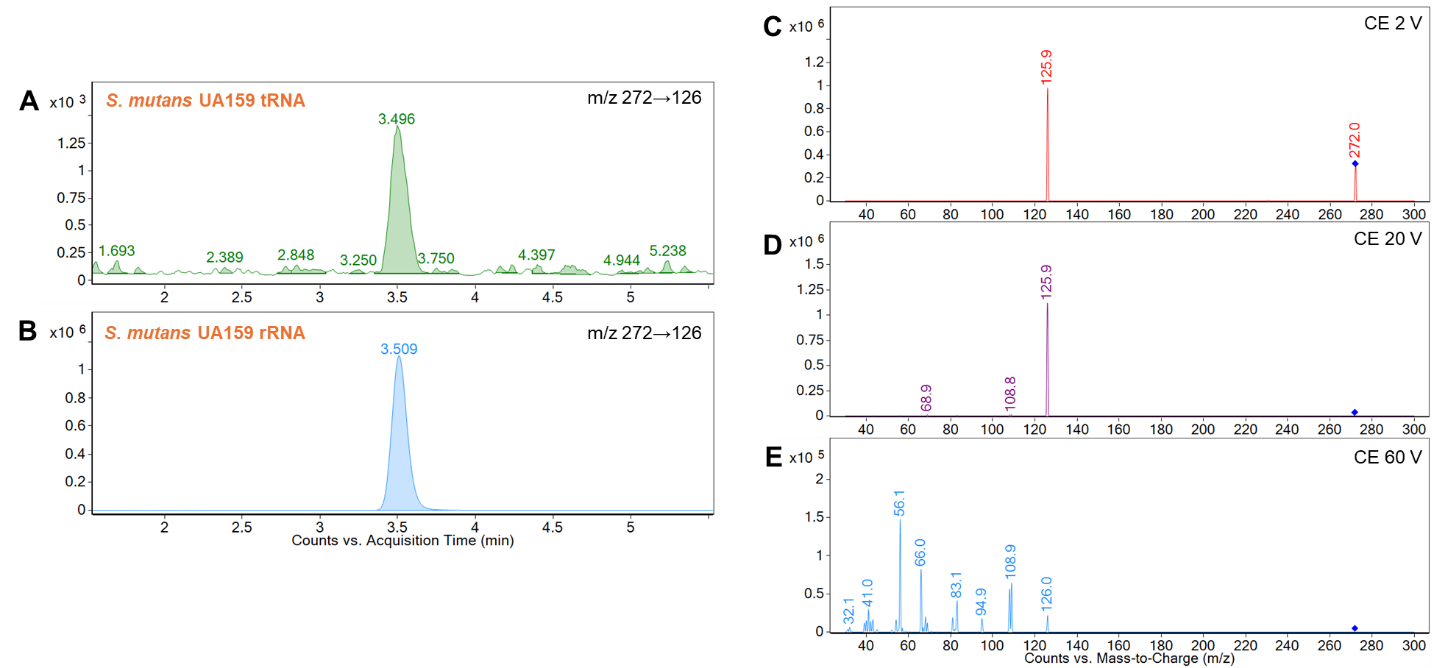

**Figure 6.** Extracted ion chromatograms obtained from LC-MS/MS analysis of (A) tRNA and (B) rRNA hydrolysate of *S. mutans* UA159 using a Waters ACQUITY UPLC BEH C18 column (50 × 2.1 mm i.d., 1.7 µm), mobile phases, LC gradient detailed in the **Methods** section. (C-E) MS/MS spectra obtained from collision-induced dissociation of the ion at *m/z* 272 in the rRNA hydrolysate using collision energies of 2 V, 20 V, and 60 V, respectively.
